# Conformational changes induced in ubiquitin by circular protein-DNA chimeras

**DOI:** 10.64898/2026.07.05.736590

**Authors:** Soumendu Boral, Michael D. Schnebly, Darartu M. Gamada, Kevin H. Gardner, Doeke R. Hekstra

**Author notes:** Corresponding authors: Kevin Gardner, CUNY Advanced Science Research Center, 85 St. Nicholas Terrace, New York, NY 10031.; Doeke Hekstra, Department of Molecular and Cellular Biology, 52 Oxford St, Cambridge, MA 02138. These authors contributed equally (Soumendu Boral, Michael D. Schnebly).

## Abstract

Proteins are dynamic molecular machines that change shape in response to physical and chemical perturbations. Although single-molecule force spectroscopy provides precise information about the stretching of proteins in response to tunable forces, it does so without structural detail. Circular protein-DNA chimeras, with DNA attached to pairs of surface sites, have been introduced as an alternative way to tunably apply forces to proteins. Intriguingly, these chimeras should be tractable for atomic-level study by nuclear magnetic resonance (NMR) spectroscopy and other structural methods. Here, we describe the NMR-scale synthesis of circular chimeras of single-and double-stranded DNA with ubiquitin, an essential component of many cellular pathways. We designed these chimeras to probe a two-residue retraction of ubiquitin’s C-terminal β5 strand, normally triggered by phosphorylation of serine 65 during initiation of mitophagy. We probed the resulting conformational changes by NMR and found that the attachment of a single strand of DNA suffices to alter this conformational equilibrium. A control bearing two separate short single DNA strands recapitulated much of the circular chimera’s NMR properties, supporting a dominant role for local protein-DNA interactions rather than spring-like action by single- or double-stranded DNA. These results provide a necessary benchmark for future studies using DNA springs to probe the functional dynamics of proteins.

**Significance statement:** Ligands, post-translational modifications, and mechanical inputs reshape proteins through forces that propagate across their structures. The underlying mechanical properties of proteins mediating these changes are rarely accessible with atomic-level detail. By producing NMR-scale circular protein-DNA chimeras, we provide a route to examining how defined physical perturbations alter protein conformational landscapes. The results establish both the promise of this strategy and the need to account for local DNA-protein interactions.

## Introduction

Proteins cycle through functional and conformational states as they catalyze reactions, bind ligands, transduce signals, and as these roles are allosterically modulated by other molecules. In each case, they respond to physical or thermodynamic forces. Despite rapid advances in structure determination and prediction, few means are available to quantitatively study protein conformational dynamics in response to physical perturbations. Nuclear magnetic resonance (NMR) spectroscopy provides one of the most direct and sensitive means to study protein conformation and dynamics, with a combination of experiments yielding atomic-level information on psec-sec interconversions, even those involving rarely-populated states (Kay, 2016). Nevertheless, techniques are lacking to combine NMR with the direct application of external forces such as those applied in single-molecule force spectroscopy (Bustamante and Yan, 2022).

Intriguingly, work by Zocchi and others has established that double-stranded DNA (dsDNA), covalently attached to a pair of sites on a protein surface, can be used to control the turnover (Tseng and Zocchi, 2013; Joseph et al., 2014) and specificity (Gokulu and Banta, 2024) of enzymes in a titratable manner. The ability of such “DNA springs” to perturb protein physics primarily arises from the fairly stiff nature of the dsDNA double helix—its persistence length of about 50 nm is much larger than the diameter of most proteins (Guilbaud et al., 2019), resulting in bending when attached to a protein surface at both ends and an outward force on the protein at the attachment sites. As a result of extensive calibration and theoretical modeling, these forces are well understood (Zocchi, 2018).

Here, we take the first steps towards systematic application of external forces using DNA springs combined with structural observation by NMR. Importantly, attachment of DNA to a protein may also lead to specific molecular interactions that complicate the naïve concept of dsDNA springs as otherwise-inert physical springs, and of ssDNA as a floppy and inert control. To establish a baseline for systematic perturbation studies, we here studied the effect of DNA spring attachment on ubiquitin (Ub), a central model system in the establishment of NMR methods (Driscoll et al., 2025). Notably, we attach ssDNA and dsDNA springs to surface sites across the C-terminal β5 strand. This strand includes residue S65, a substrate of PINK-1 kinase. Mutations in PINK-1 cause Autosomal-recessive juvenile Parkinsonism. Gladkova et al. established the existence of an excited state in which this C-terminal β strand retracts in a −2 register shift to expose S65 at the surface (Gladkova et al., 2017). This retracted conformation is likely essential for phosphorylation by PINK-1, with failure to do so disrupting the clearance of defective mitochondria (Schubert et al., 2017).

We first established a protocol for the synthesis of circular DNA-Ub chimeras with sufficient yield and purity to enable solution NMR studies. We found that attachment of ssDNA, at positions 9 and 63, strongly affected the central β-sheet of ubiquitin and shifted the C-terminal β5 strand’s conformational equilibrium towards a retracted state.

By comparison, addition of a complementary strand to form a dsDNA spring has modest effects, emphasizing that local interactions, possibly complemented by the non-trivial persistence length of ssDNA, can substantially influence the outcomes of using DNA springs to probe the structural response of proteins to applied forces.

## Results

### Protein Biochemistry

#### Synthesis of circular protein-DNA chimeras

Perturbation of the conformational ensemble of a protein in solution requires, minimally, the application of a pair of forces. Ubiquitin provides an interesting model system for direct perturbation studies because of its widespread biological importance and the presence of a strand-retraction equilibrium implicated in Parkinson’s disease (Wauer et al., 2015; Gladkova et al., 2017). To apply forces to pairs of sites on the surface of ubiquitin, we needed to develop a protocol to produce circular protein-DNA chimeras with sufficient scale and purity for solution NMR studies. Our approach builds on prior work on protein-DNA chimeras (Mukhortava and Schlierf, 2016; Tseng et al., 2021), and is shown schematically in **Figure 1** with a detailed description provided in the **Methods**. Central to this approach is the attachment of DNA, through heterobifunctional linkers, to surface cysteine residues. Since (human) ubiquitin does not natively contain cysteines, we introduce these by site-directed mutagenesis (here T9C and K63C). All studies reported here were performed in the background of the F45W mutation, a well-characterized mutant which increases UV absorbance nearly 5-fold with little effect on its biophysical properties (Khorasanizadeh et al., 1993).

**Figure 1:**
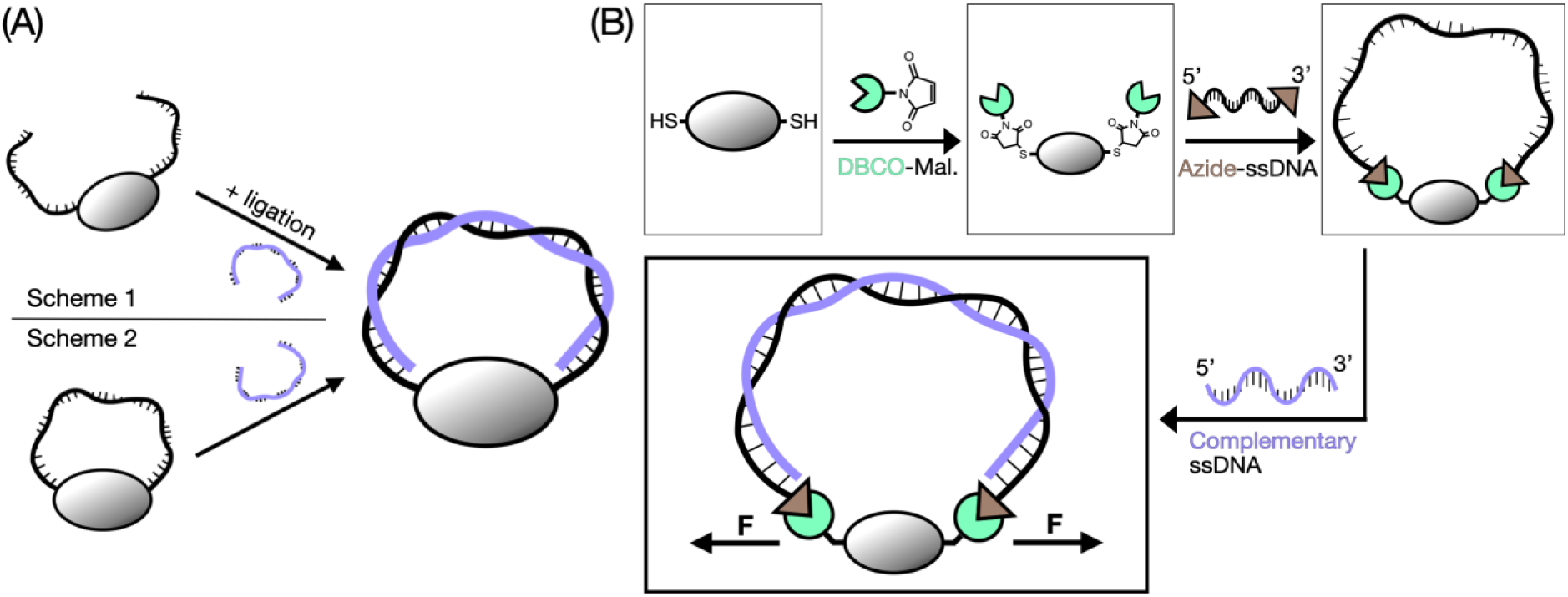
Protein-DNA spring synthesis. (A) Two schemes for creating circular protein-DNA chimeras. Figure adapted from (Wang et al., 2009). (B) Synthesis chemistry and workflow for Scheme 2. The protein of interest, containing two surface cysteines, is conjugated to azide-modified ssDNA using a DBCO-maleimide cross-linker. Upon addition, the complementary strand hybridizes with the covalently attached strand to form a spring-like, bent double-helix.

Zocchi and colleagues (Wang et al., 2009) outlined two strategies for making circular protein-DNA chimeras (**Fig 1A)**. In the first, two kinds of short oligos are added, one modified with a cross-linkable group at the 5’ end and the other modified at the 3’ end. After attachment, a complementary strand is then added, followed by ligation of the free ends of the first two strands. This strategy is robust to incomplete modification of oligos. A drawback is the formation of dimeric chimeras, which are favored before ligation (Wang and Zocchi, 2009). As a consequence, this protocol does not scale well to the concentrations needed for NMR which requires ∼10^5^⊆ more material. As an alternative, they described a second approach using an initial oligo modified with cross-linkable groups at both ends but dismissed this option due to the difficulty avoiding linear species due to missing cross-linkable groups (Wang et al., 2009).

Here, we pursued this second approach, starting from a single DNA strand modified at both 5’ and 3’ ends with azide groups (Integrated DNA Technologies, Inc.), made possible by improved oligo modification and an effective purification protocol described below. We covalently attached this single-stranded DNA to surface cysteine residues using the heterobifunctional crosslinker dibenzocyclooctyne (DBCO)-maleimide (Vector Labs), attaching to azides and cysteines via its DBCO and maleimide moieties, respectively (**Fig 1B**). The chemistry is inspired by (Mukhortava and Schlierf, 2016), but distinct in that their synthesis required only single-modified DNA.

The bivalent nature of both the protein and DNA leads to formation of a variety of chimeric products (**Fig 2; Fig S1**). To identify the SDS-PAGE band associated with each product, we first identified products of a synthesis reaction using single-cysteine (K63C) ubiquitin, for which the number of possible products is necessarily limited. Using this comparison, we established that the circular ssDNA-Ub chimera runs at a slightly smaller apparent molecular weight than the linear chimera (**Fig 2**). As expected, we found the optimal reactant ratio for producing circular ssDNA-Ub to be 1:1 (**Fig S2**).

**Figure 2:**
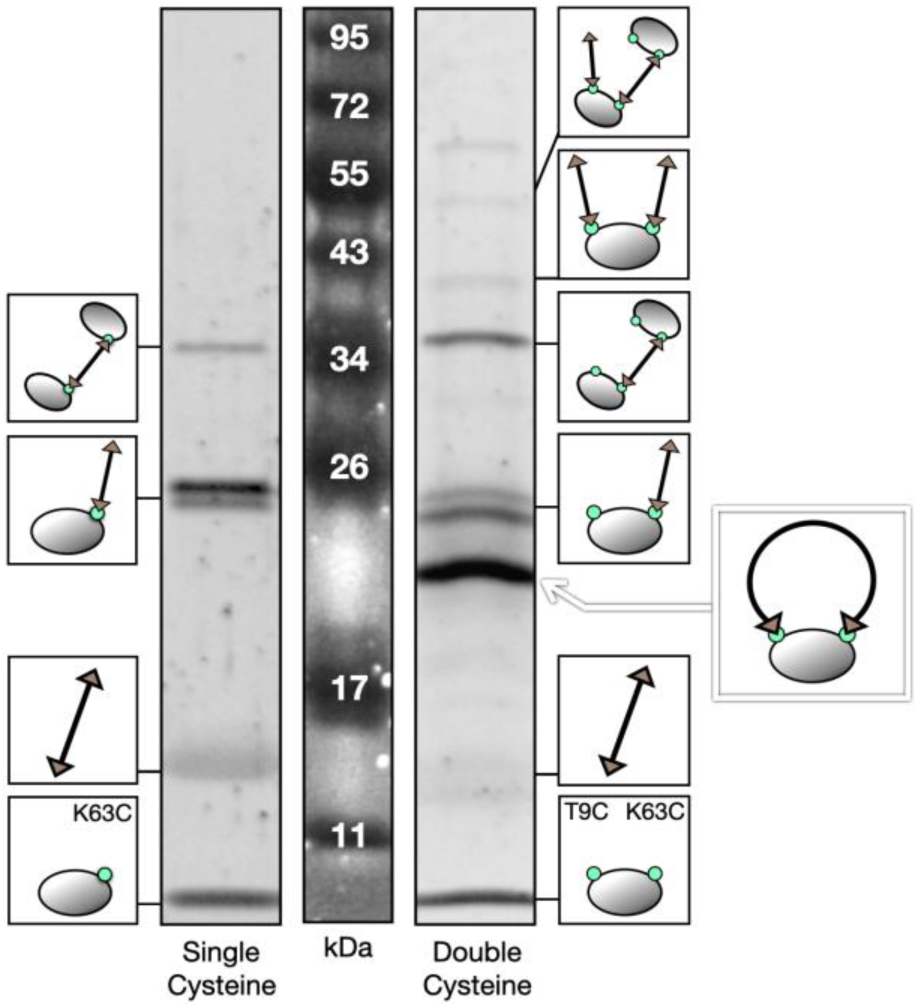
Identification of ssDNA-Ub synthesis products in SDS-PAGE. Single-cysteine controls prove useful for initial band identification due to the reduced number of potential products. While products tend to run near their molecular weight, isomeric products may run at different sizes, as is the case with linear ssDNA-Ub (∼25 kD) vs. circular ssDNA-Ub (at ∼20 kD). Note that DNA is stained slightly by the protein stain, SYPRO Ruby.

#### Purification of circular ssDNA-Ub chimeras

We purified circular ssDNA-Ub in two steps. First, we used size exclusion chromatography to separate products with large differences in molecular weight (**Fig S3**). Next, we “cleaned up” remaining non-circular product (**Fig 3A**). Most non-circular products contain unreacted moieties. Using this to our advantage, we covalently captured these groups using agarose beads modified to contain their reactive counterparts (**Fig 3A**). Specifically, maleimide-agarose (Cube BioTech) binds and sequesters products containing unreacted cysteines while DBCO-agarose (Vector Labs) does the same for unreacted azides. Use of azide-agarose did not have a substantial effect on purity, suggesting incomplete modification of starting oligos with azide groups as a dominant cause of incomplete product formation.

**Figure 3:**
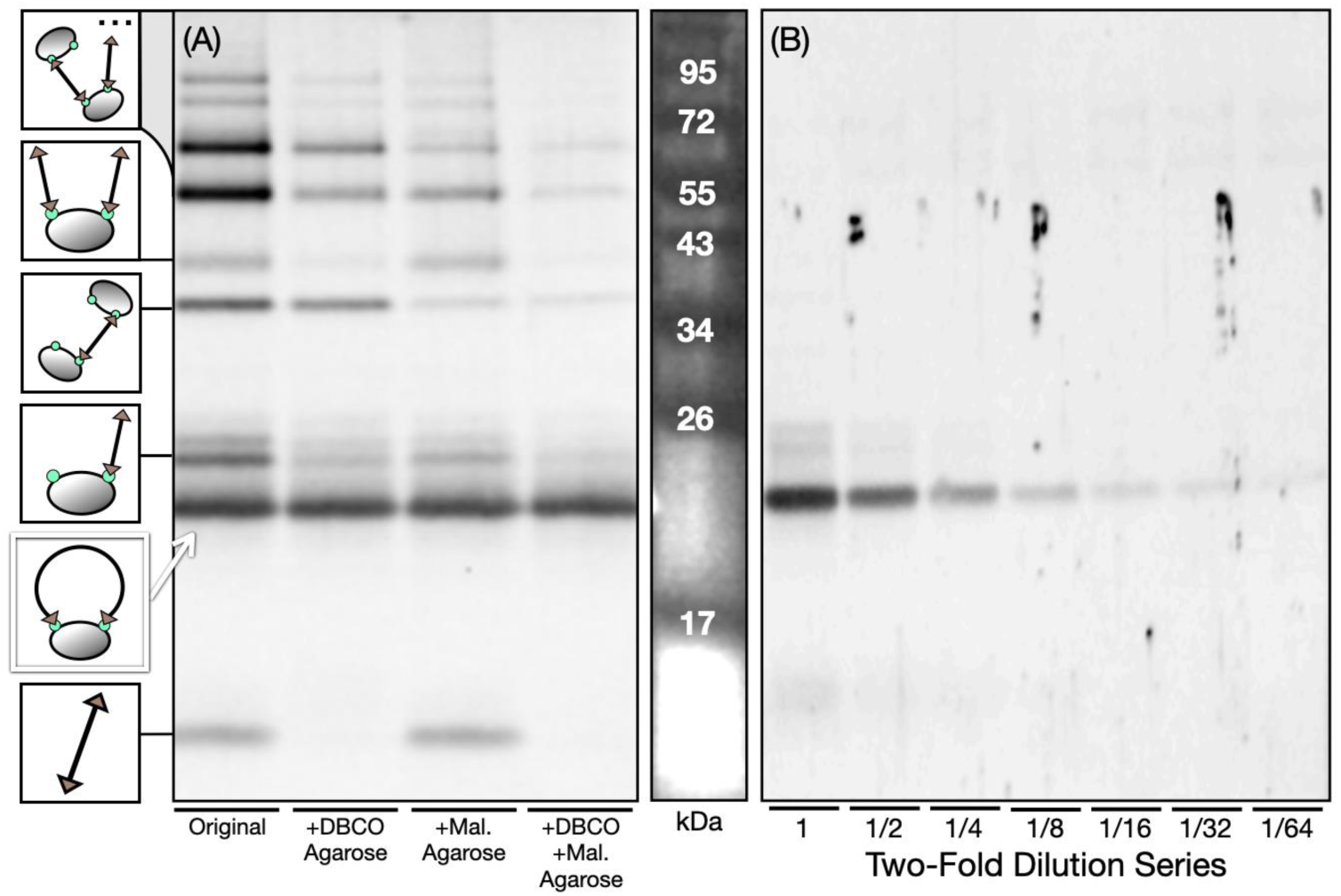
Purification procedure and purity assay for circular Protein-ssDNA. (A) Non-circular products contain reactive subgroups. Each can be sequestered by its counterpart in functionalized agarose beads. For example, DBCO-agarose conjugates with products containing unreacted azides, removing them from solution. Similarly, maleimide-agarose sequesters species with free cysteines. (B) Estimating purity of circular product. After purification, we tend to be left with trace amounts of linear isomer. Here, the circular product disappears five two-fold dilutions after the linear product disappears, indicating it is about 32 times more abundant.

For synthesis and purification of the circular U-[^13^C,^15^N] DNA-Ub sample used here, 1.5 μmol of apo-Ub were converted to 1.5 μmol DBCO-Ub, then combined with 1.5 μmol of azide-ssDNA in SEC buffer to create a 500 mL, 3 µM 1:1 mixture. After concentration and purification, the sample contained 390 nmol of circular ssDNA-Ub for a synthesis yield of ∼25%. The resulting samples contained circular U-[^13^C,^15^N] protein-DNA chimeras at a quantity (500 µL of 150-300 µM) and quality (>95% pure; **Fig 3B**) sufficient for NMR studies.

#### Formation of dsDNA-Ub chimeras by addition of complementary strand

To form dsDNA-Ub chimeras, we combined purified circular ssDNA-Ub, as just described, with its complementary ssDNA strand, hybridizing to form a bent double-helix. Linear and circular dsDNA-Ub chimeras migrated at different speeds in SDS-PAGE than would be expected based on the complementary strand’s added mass (**Fig 4; Fig S4**), consistent with previous findings (Gaillard and Strauss, 2015; Waszkiewicz et al., 2023). We found that a 1.2x molar excess of ssDNA was sufficient to produce a complete band shift from Ub-ssDNA to Ub-dsDNA chimera (**Fig 4**).

**Figure 4:**
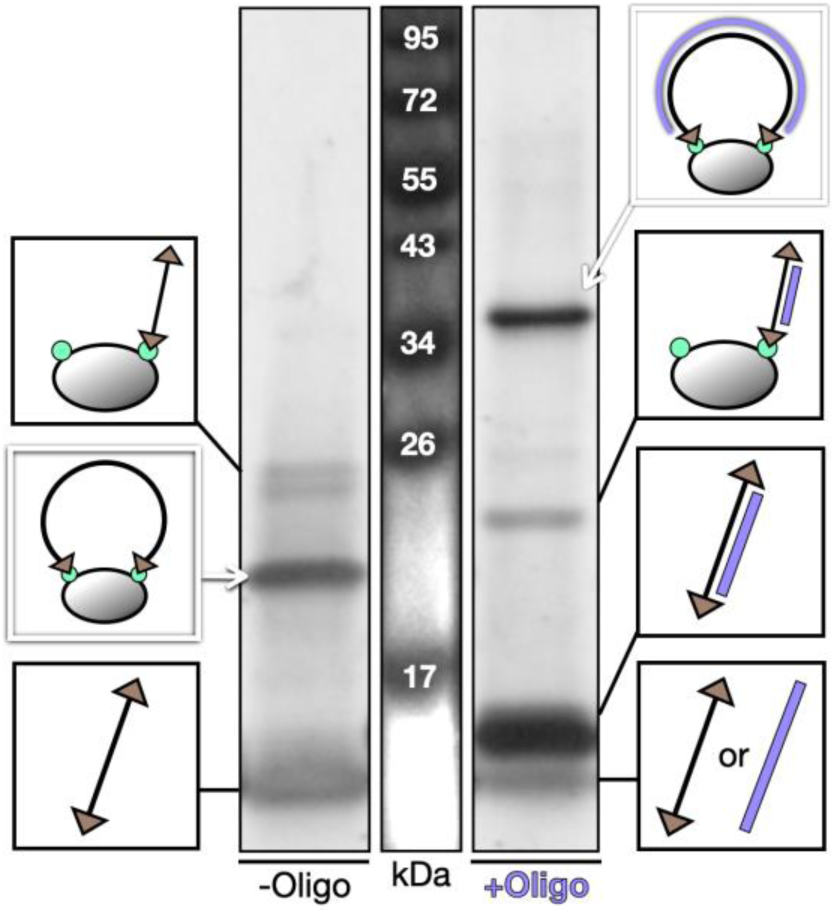
Complementary DNA strand hybridization confirmed by band shift in SDS-PAGE. Hybridization with complementary strand, added here at a 1.2x molar excess, causes band shifts that may be unintuitive. Increased rigidity of the double-helix alters the mechanics of product passage through the polyacrylamide gel matrix. Linear products tend to move faster, while circular products move slower. SDS does not denature dsDNA. For further evidence of these band identifications, see **Figure S4.**

### Nuclear Magnetic Resonance (NMR) spectroscopy

#### NMR chemical shift changes in Ub due to covalent attachment of DNA

To maximally perturb the β5 strand, we incorporated DNA attachment sites near its base (K63C) and at a site diametrically across the protein and not part of the β5 strand (T9C) (**Fig 5C**). As expected from the high structural stability of ubiquitin, the ^15^N/^1^H HSQC spectrum of apo-Ub showed well-dispersed peaks indicating that the mutations introduced for DNA tethering do not substantially perturb tertiary structure (**Fig 5A**; **Fig S5A**). Covalent attachment of 50-nt single-stranded DNA led to substantive chemical shift changes (**Fig 5A; Fig S5B**), suggestive of structural changes within the protein, along with peak broadening as anticipated from the increased molecular weight of the complex (adding 17 kD of DNA yields 25.5 kD total mass for ssDNA-Ub).

**Figure 5:**
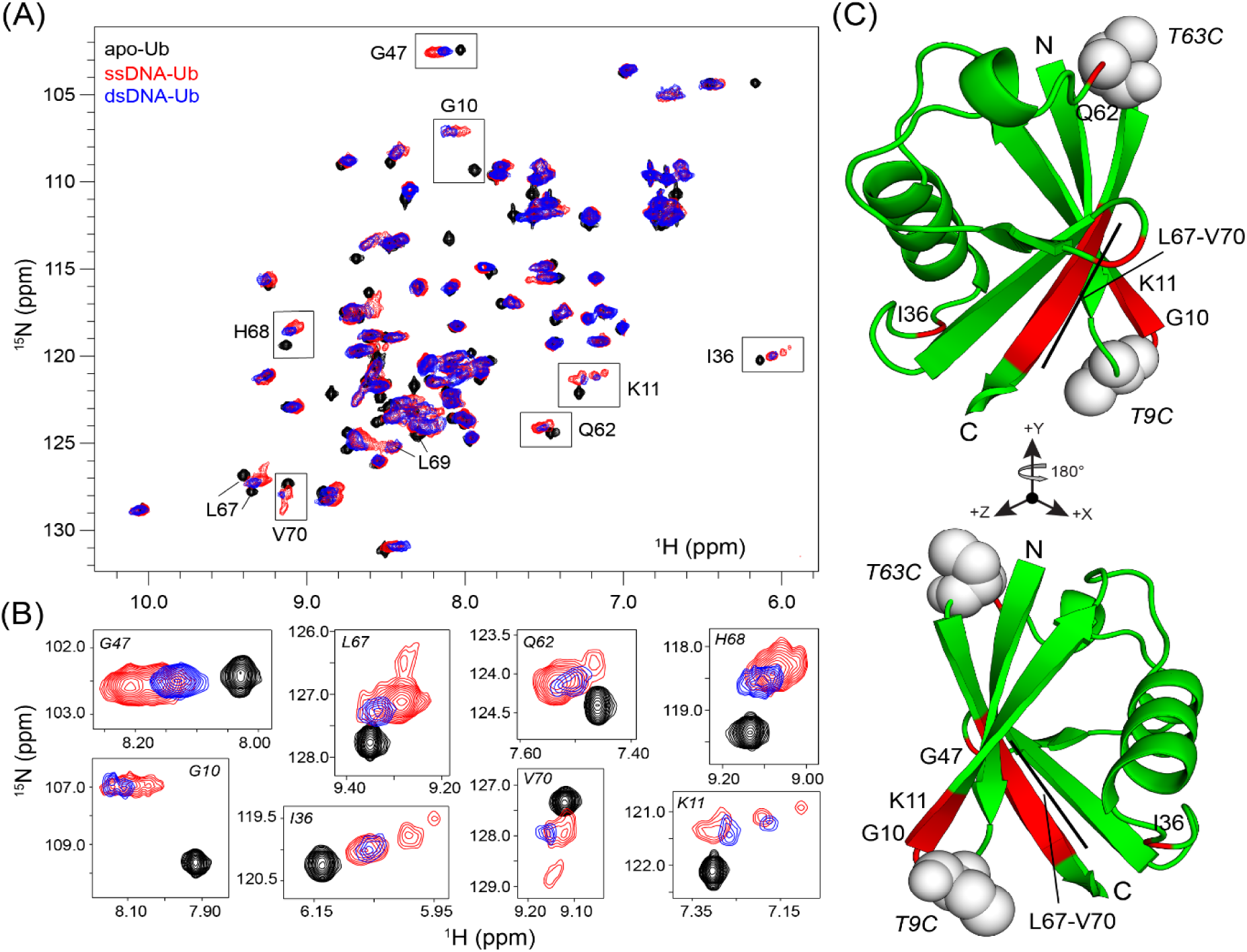
DNA tethering induces conformational changes in Ub without disrupting overall tertiary fold. (A) Overlay of ^15^N/^1^H HSQC spectra of apo-Ub (black), ssDNA-Ub (red), and dsDNA-Ub (blue) at 40°C, pH 5.8. Residues showing multiple conformations in the presence of ssDNA are indicated with boxes. (B) Overlay of ^15^N/^1^H HSQC spectra for selected resonances, using the same spectra as (A). (C) Residues showing multiple conformations are mapped with red color onto the wildtype Ub structure (PDB 1UBQ). The DNA tethering sites (T9C and K63C) are shown as spheres; specific residues and N/C-termini are labeled. Structures are generated using PyMOL 3.1.8.

Addition of the complementary DNA strand led to further broadening, as expected by increased molecular weight to a total of 42.5 kD for the dsDNA-Ub complex (**Fig 5A**; **Fig 5B**; **Fig S5C**). We confirmed that the newly added complementary strand formed Watson-Crick hydrogen bonds with the existing covalently-attached strand by 1D ^1^H NMR, focusing on the ^1^H 12-14 ppm DNA imino proton region. While only two weak peaks were apparent in this spectral region for ssDNA-Ub, perhaps due to some intramolecular hydrogen bonding, we observed a large number of peaks for dsDNA-Ub consistent with Watson-Crick base pairing and hydrogen bonding of the imino protons of guanosine and thymidine deoxynucleotides (**Fig S6A**). These imino signals remained intact across the 25-40°C range of our experiments (**Fig S6B**), confirming the stability of the DNA duplex. Notably, the addition of the complementary strand had only minor effects on the protein ^15^N/^1^H^N^ signals, suggesting minimal conformational changes of the protein components of the ssDNA-Ub and dsDNA-Ub complexes (**Fig 5**). At all points, ubiquitin clearly remained folded as demonstrated by overlays of ^15^N-^1^H HSQC and ^13^C/^1^H constant-time HSQC spectra for all three samples (**Fig 5A; Fig S5; Fig S7**).

To assess the structural changes caused within ubiquitin by covalently attaching ssDNA or dsDNA, we used standard triple resonance NMR methods (Sattler, 1999) with uniformly-^13^C,^15^N protein-labeled samples to assign the protein chemical shifts of each Ub variant. With this approach, we assigned more than 94% of backbone atoms for all the three different constructs (**Table S1**) with coverage at virtually all of 73 non-proline residues (70 backbone amides and associated ^13^Cα, ^13^Cβ, and ^13^CO for apo-Ub mutant, and 68 backbone amides and nearby ^13^C atoms for ssDNA-Ub and dsDNA-Ub adducts) (**Fig S8**).

#### Conformational and dynamics changes within Ub caused by tethering DNA

To further explore the roots of the protein chemical shift changes caused by tethering DNA, we recorded ^15^N/^1^H HSQC spectra of ^15^N-labeled ubiquitin with different covalent modifications. The addition of DBCO linkers to apo-Ub (forming DBCO-Ub) led to substantial heterogeneity at residues near the C9 and C63 tethering sites (in particular the L8 loop (residues 7-11) and at residue 62; **Fig S9**), consistent with local restrictions around the modified cysteines generating conformers in slow exchange. In addition, residues 36, 67, and 70 showed multiple conformers in slow exchange. This is plausibly due to coupling of these residues with the L8 loop (through a hydrophobic patch involving residues 8, 44, 68, and 78 (Komander and Rape, 2012)) rather than propagation of strain along the β5 strand which is not otherwise strongly affected by attachment of just the linkers.

Proceeding to tether cysteines 9 and 63 together with a 50 nt ssDNA bridge (ssDNA-Ub), we made two key observations. First, conformational heterogeneity near the tethering sites was strongly reduced (e.g., at positions 10 and 62) but not everywhere (e.g., at position 11, **Fig S9**). Secondly, residues along the β5 strand (residues 64-72 in the ground state) and the C terminus showed chemical shift changes in response to DNA attachment larger and different than those from attachment of the linker alone (**Fig S9**). Subsequent addition of equimolar quantity of complementary strand at 40°C further reduced the number of alternate conformations in Ub, often down to a single of several conformers seen in the ssDNA-Ub spectrum (**Fig 5A**; **Fig 5B; Fig S9**). Similarly, increased temperature altered the relative populations of the heterogenous peaks for any given amide (**Fig S10**).

Using the chemical shifts of backbone and Cβ atoms (^1^H^N^, ^15^N, ^13^Cα, ^13^Cβ, and ^13^CO), we more closely examined the changes in structure and dynamics among the apo-, ssDNA-, and dsDNA-Ub constructs. We started by comparing apo-Ub (T9C / F45W / K63C) to the published chemical shifts of wildtype Ub (Cornilescu et al., 1998). Our analysis of ^1^H^N^ and ^15^N chemical shift changes between these two proteins (**Fig. 6A**) showed that alterations were restricted to the immediate vicinity of the mutated residues and without substantial long-range perturbations, as intended.

**Figure 6:**
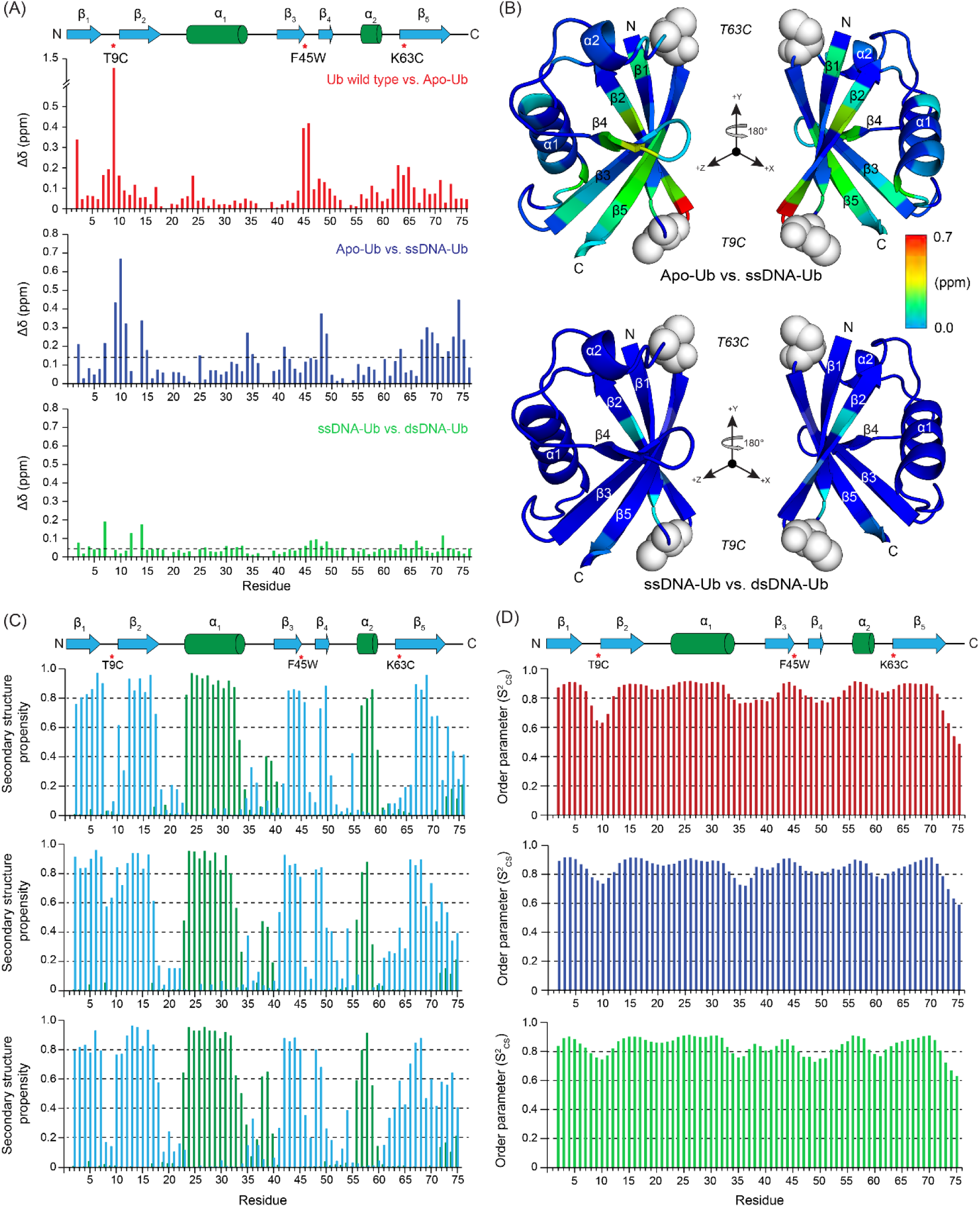
Site-specific NMR analyses reveal DNA tethering to Ub causes C-terminal conformational changes. (A) Backbone ^1^H and ^15^N chemical shift differences between wild-type and apo-Ub mutant (upper), apo-Ub mutant and ssDNA-Ub (middle), and ssDNA-Ub and dsDNA-Ub (bottom); average Dd values are indicated by dotted lines on each plot. (B) DNA-induced chemical shift perturbations are mapped onto the Ub structure (PDB: 1UBQ) with spheres denoting the DNA attachment sites. The blue-to-red color scale in ppm unit is shown on the right side. The β-strands, attachment sites, N and C-termini are indicated. Structures are generated using PyMOL 3.1.8. (C) MICS-derived backbone chemical shift-derived secondary structure propensities of different Ub variants are plotted for each residue (from top to bottom: Apo-Ub, ssDNA-Ub, dsDNA-Ub). The propensities for helices and strands are colored in green and blue, respectively. (D) MICS-derived backbone chemical shift-derived order parameters (S^2^_CS_) are plotted for each residue. For all plots, the apo-Ub secondary structure and sites of mutations are indicated schematically at the top.

To examine the effects of covalently attaching DNA to Ub, we compared chemical shifts between the apo-, ssDNA-, and dsDNA-Ub samples (**Fig 6A**). The ^1^H^N^ and ^15^N chemical shift differences we observed between these sets established that ssDNA binding affected sites throughout the protein. Mapping these shift changes onto the crystal structure (**Fig 6B**), we observed many of the largest differences on the regions around the attachment sites and within the β5 strand. Taken together with the observed impact of linker attachment itself (above and **Fig S9**), these data indicate that attachment of single-stranded DNA to apo-Ub triggers conformational change extending through the H-bonding network of ubiquitin’s curved β-sheet. In contrast to this substantive change, the subsequent annealing of the complementary DNA strand to convert ssDNA-Ub to dsDNA-Ub had only modest effects.

We further used this chemical shift information to calculate secondary structure propensities and order parameter values for all three constructs using MICS (Shen and Bax, 2012). Secondary structure calculations for the ssDNA-Ub and dsDNA-Ub species confirmed that their modifications did not affect the overall α/β topology of the apo-Ub mutant, consistent with the limited changes in ^15^N/^1^H^N^ chemical shifts among these samples and modest applied perturbations (**Fig 6A, 6C**). The most substantial changes were seen in the C-terminal region of the protein, with increased β-strand propensities in both the ssDNA-Ub and dsDNA-Ub complexes compared to the apo-Ub mutant (**Fig 6C**), consistent with the −2 register shift of β5 inherent to Ub (*i.e*. increased β propensity in residues 71-75).

These observations strongly suggested that DNA tethering increases the probability of the C-terminal residues to be included in the β-sheet. Importantly, the attachment of ssDNA sufficed to accomplish this. Given the short persistence length of ssDNA this suggested that local interactions with the protein, more than spring-like action of the DNA, mediate its effect on ubiquitin’s conformational equilibrium. To further test this, we also constructed a chimera in which we tethered two separate 12-nt ssDNA strands to ubiquitin (**Fig S9**). Consistent with the local interaction hypothesis, this ssDNA-Ub(s) chimera displayed a ^15^N-^1^H HSQC spectrum strongly resembling that of the full circular ssDNA-Ub chimera.

#### Identification of Ubiquitin C-terminal retracted conformation upon DNA tethering and its stabilization

Previous NMR studies of ubiquitin have identified a high-energy alternative conformation with a −2 shift in the register of the β5 strand compared to the native structure (Wauer et al., 2015). This “C-terminal retracted” (CR) conformation of Ub was initially identified as a rarely populated species for wild-type Ub (<1% at 45°C) and shown to be stabilized by PINK1 kinase and was subsequently found to be more highly populated upon phosphorylation of β5-residue S65 (∼30% at 25°C) (Gladkova et al., 2017). Following the increase in β-strand propensity and order parameter values in the C-terminal β5-strand after tethering ssDNA (**Fig 6B, 6C**), we suspected that this covalent modification might promote formation of the CR conformation. To test this concept, we obtained ^1^H^N^-^1^H^N^ NOEs from a 125 ms mixing time ^15^N-edited NOESY experiment recorded on a 300 µM ssDNA-Ub sample. Such NOESY-type spectra provide distance information between protons separated by up to ca. 5-6 Å, which are particularly useful to establish the relative register of adjacent strands within a β-sheet.

Strikingly, the ^15^N-edited NOESY data revealed that ^1^H^N^-^1^H^N^ NOEs between the C-terminal β5 strand and the adjacent β1 and β3 strands are perturbed by ssDNA attachment (**Fig 7**), including NOEs consistent with distances up to ∼5.5 Å from either wild type (PDB: 1UBQ; (Vijay-kumar et al., 1987)) or the −2 slipped conformations of Ub (PDB: 5OXI; (Gladkova et al., 2017)) (**Table S2**). These observations strongly suggest that both conformers exist in the sample. Notably, three observed ^1^H^N^-^1^H^N^ NOEs (Q40-L73, R42-L69, S65-I3) did not have distances under ∼5 Å in either structure, suggesting either potential conformational dynamics not well represented in the crystal structures or spin diffusion during the 125 ms mixing time. We favor the former interpretation for most of these NOEs; for example, the L73 residue is not well-defined in 1UBQ, opening the possibility that a shorter distance may be populated, biasing the NOE peak intensities during conformational exchange (Kim and Prestegard, 1989). Together with chemical shift-derived information, these ^1^H^N^-^1^H^N^ NOEs strongly suggest that covalent tethering of ssDNA to sites 9 and 63 stabilizes the C-terminal β5 slipped conformation sufficiently to impact several NMR indicators of the Ub structure and dynamics.

**Figure 7:**
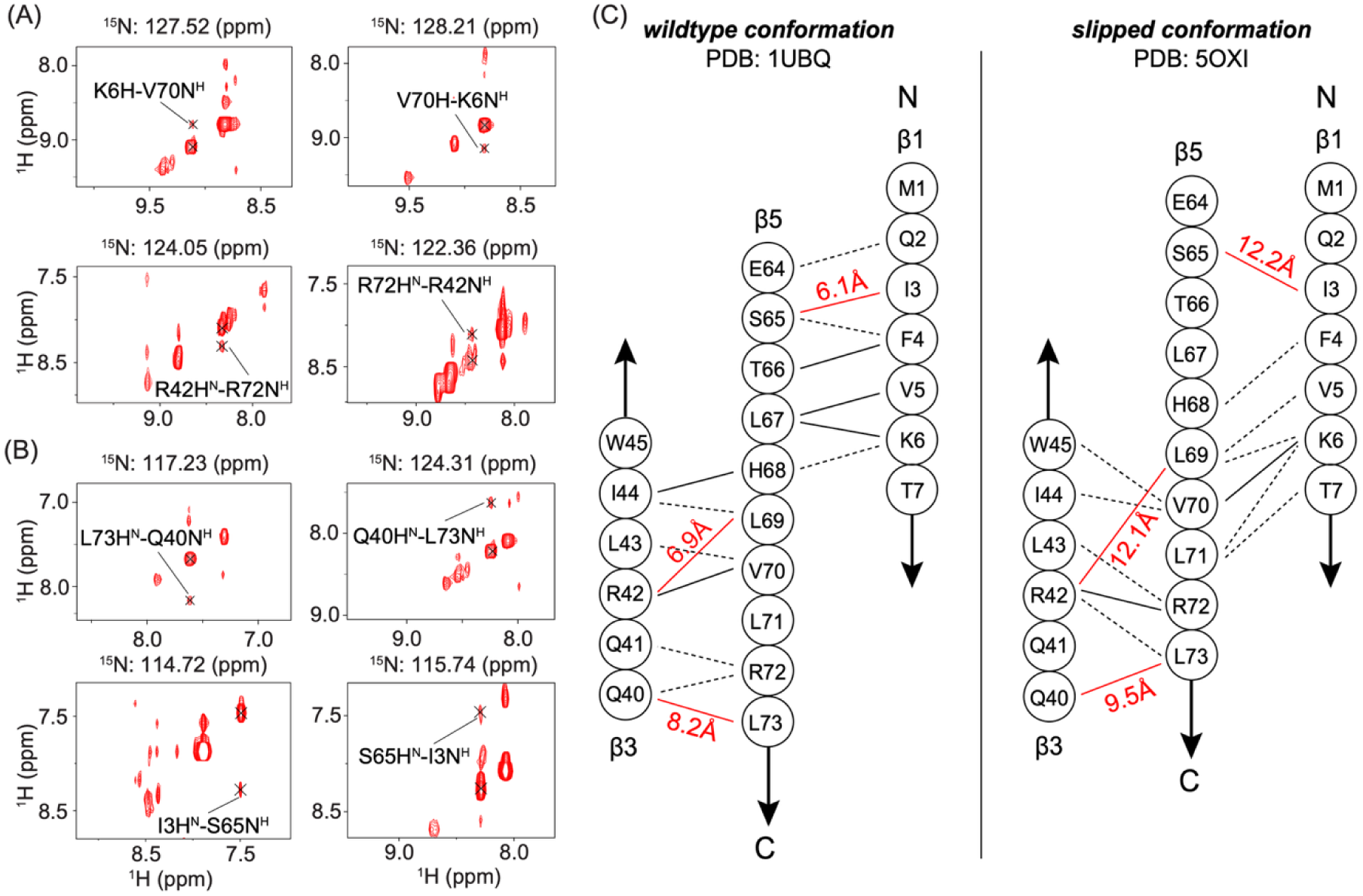
NOESY data for ssDNA-Ub support a DNA-induced increase in slipped β-sheet population of Ub domain with covalently-tethered single stranded DNA. (A) Representative ^15^N-edited NOESY strip plots showing two pairs of ^1^H^N^-^1^H^N^ NOEs (K6-V70, R42-R72) uniquely < 5 Å in the slipped conformation. (B) Representative ^15^N-edited NOESY strip plots showing two pairs of ^1^H^N^-^1^H^N^ NOEs (Q40-L73, I3-S65) suggestive of conformational dynamics involving residues at either end of the β5 strand. (C) Schematic representations of the wild-type and slipped Ub conformations, with ^1^H^N^-^1^H^N^ NOEs between the β5-β1 and β5-β3 strands depicted. Predicted NOEs (^1^H^N^-^1^H^N^ distances ≤ 5.5 Å) are shown as black dotted lines; predicted and observed ones are shown as black solid lines, and newly observed NOEs are shown as solid red lines (labeled with ^1^H^N^-^1^H^N^ distances from individual PDBs). Other distances are listed in **Table S2**.

## Discussion

Our work demonstrates the ability to generate tethered DNA-protein samples at scale suitable for NMR studies, evidence for their utility to influence a pre-existing conformational equilibrium within a protein, and potential for careful study of the structural basis of such effects.

Covalent attachment of single-stranded DNA to ubiquitin at sites T9C and K63C induced significant chemical shift changes and peak broadening in NMR spectra, indicating structural perturbations without disrupting the protein’s overall fold (**Fig. 5**). The subsequent annealing of a complementary DNA strand to form double-stranded DNA further increased molecular weight and broadened peaks, but had minimal additional impact on Ub’s chemical shifts, confirming duplex formation via Watson-Crick hydrogen bonding and stable imino proton signals across temperatures (**Fig. S6**). Detailed NMR analyses confirmed that Ub remained folded across all constructs, with over 94% backbone atom assignments achieved, confirming structural integrity. The crosslinkers introduced conformational heterogeneity, particularly near the attachment sites and the β5 strand, with multiple chemical shifts suggesting slow exchange among distinct conformers. This heterogeneity was reduced upon duplex formation or elevated temperature, suggesting dynamic equilibrium among Ub conformers influenced by DNA interactions.

The conformational changes in ubiquitin induced by single-stranded DNA attachment resemble the alternative structure previously described by (Gladkova et al., 2017), as supported by NMR-based chemical shift changes and measurements of secondary structure propensities, backbone dynamics, and ^1^H^N^-^1^H^N^ NOEs. We propose that the subsequent addition of the complementary strand helps stabilize this non-native conformation. ^1^H^N^-^1^H^N^ NOE analysis is particularly informative as it reveals patterns consistent with both the ssDNA-Ub complex and the −2 slipped β5 strand register, suggesting coexistence rather than a full conformational transition. Although we lack quantitative data to determine the relative populations of these states, the evidence points to the presence of both conformers within the sample. These species likely interconvert on the timescale captured by NOE measurements, reflecting a dynamic equilibrium influenced by DNA tethering.

Taking a general view, DNA springs provide a potentially potent means to quantitatively perturb the soft modes of proteins often associated with functional conformational transitions. Particularly interesting is the potential to tune the applied forces by controlling the length of the primary tethered DNA strand and the extent of hybridization with a complementary DNA strand. To lay the foundations for such studies, it is critical to address basic questions regarding use of DNA springs. Using ubiquitin as a model system, we here performed such a study and found a strikingly strong effect from just attachment of the primary ssDNA strand. A control experiment tethering two separate short ssDNA strands to Ub ruled out a spring-like mechanism as a dominant factor. Instead, the results point to the importance of direct local interactions between the DNA and protein, although a role for ssDNA stiffness at low salt concentration (Chen et al., 2012) cannot be excluded. The limited effect of adding complementary DNA further suggests that the β5 strand retraction equilibrium is neither a soft mode along which the protein can readily be deformed by picoNewton forces due to bending of dsDNA, nor a discrete equilibrium strongly biased by the applied force. By way of illustrative calculations, at the end-to-end distance used here (∼2.5 nm), a 50-bp dsDNA spring is prone to buckling, resulting, effectively, in a molecular tweezer with ∼8 nm arms and a maximum torque of about 30 pN nm, or ∼4 pN of total force (Zocchi, 2018). For compact, well-folded proteins, spring constants are commonly in the range of 100 *kT*/nm^2^ (Zocchi, 2018), leading at this applied force to continuous displacement of about 0.1 Å. Our results are therefore compatible with limited deformation of a well-folded protein. Considering the discrete β5 strand retraction equilibrium, position 63 would be expected to shift by ∼2.8 Å along the direction of applied force, comparing PDB IDs 1UBQ and 5OXH. With a 4 pN applied force, this would result in a ∼1.3⊆ increase in the population of the minor state, which may remain hard to observe given the low occupancy of the retracted state.

Our study’s limitations provide a starting point for future works. For instance, structural effects are likely a function of tethering sites, DNA composition and length, and cation species and concentration. In addition, it may be possible to map in more detail the interactions between the protein and ssDNA and the nature of the excited states induced by DNA tethering.

In summary, we have established that ubiquitin-DNA chimeras are tractable by NMR, whether using single-stranded or double-stranded DNA. This indicates that DNA chimeras of compact single- or two-domain proteins may be readily tractable by NMR, potentially opening up these proteins to systematic study using DNA springs.

## Materials and Methods

### Protein Biochemistry

#### Ubiquitin expression and purification

The human ubiquitin mutants F45W, F45W/K63C, and F45W/T9C/K63C were created by site-directed mutagenesis of an ampicillin-marked pET15 plasmid (AddGene: #12647). Each plasmid was transformed into chemically-competent BL21(DE3) *E. coli* (New England Biolabs), plated on ampicillin-selective LB agar, and incubated overnight at 37°C. 20 mL starter cultures were inoculated with ∼20 freshly grown colonies and grown to saturation overnight, shaking at 37°C. All 20 mL was transferred to 2 L of pre-warmed media and incubated shaking at 230 RPM, 37°C. When culture reached an OD_600_ of 0.9 (4-6 h), expression was induced by adding 400 µM IPTG. After an additional 4 hours, cells were harvested via centrifugation at 4200 xg RCF for 20 minutes at 4°C. All cultures were grown in M9 minimal media containing ampicillin and natural abundance or isotopically-enriched sources of nitrogen (^15^N NH_4_Cl) and carbon (U-^13^C glucose), both from Cambridge Isotope Labs.

To purify ubiquitin, we used a tag-free, cation exchange-based purification method (Pickart and Raasi, 2005). Cell pellets were resuspended to 50 mL in Lysis Buffer (50 mM Tris pH 7.6, 1 mM TCEP, 0.4 mg/mL lysozyme, 0.02% v/v NP-40, Roche “cOmplete” Protease Inhibitor Tablet), then incubated at 4°C, stirring for 30 minutes. Next, DNA was digested by adding DNase I to 20 µg/mL and MgCl_2_ to 10 mM and stirring for 20 minutes at 20°C. After centrifugation at 32,000 *g* RCF at 4°C for 30 minutes, the supernatant was collected and transferred to a small beaker and 0.35 mL of 70% perchloric acid was added, followed by stirring at 4°C for 10 minutes. After another round of centrifugation (same conditions), the supernatant was collected and immediately dialyzed into Cation Exchange Buffer A (50 mM Ammonium Acetate pH 4.5, 1 mM EDTA, 1 mM TCEP) in two consecutive 4-hour rounds, each with 4 L of fresh buffer at 4°C. Cation exchange was performed on an ÄKTA Pure chromatography system with a HiPrep SP FF 16/10 cation exchange column (Cytiva) using a linear gradient from 100% Buffer A to 100% Buffer B (50 mM Ammonium Acetate pH 4.5, 500 mM NaCl, 1 mM EDTA, 1 mM TCEP) over 400 mL (40 column volumes). Fractions containing ubiquitin were pooled, concentrated and then further purified by size exclusion chromatography (SEC) using a HiLoad Superdex 75 16/600 PG column (Cytiva) with SEC Buffer (30 mM PBS pH 7.4, 150 mM NaCl).

#### ssDNA-Ubiquitin chimera synthesis and purification

To synthesize circular ssDNA-Ub chimeras, the heterobifunctional crosslinker DBCO-maleimide (Vector Labs) was used to covalently attach ubiquitin containing two surface cysteines (F45WT9CK63C) to ssDNA modified to contain azides on both its 5’ and 3’ end (“azide-ssDNA”; 50 nucleotides; /5AzideN/TGGTGGAAGCTTG-TGTGGTGGTGGACATGGAGACTCAGGCGCAGTATGGC/3AzideN/; Integrated DNA Technologies). All chemistry was done in SEC Buffer.

Before mixing with azide-ssDNA, DBCO-maleimide was attached to ubiquitin as follows: to reduce disulfide bonds, purified ubiquitin was incubated in SEC Buffer with a 20-fold molar excess of TCEP to cysteines for 30 minutes at room temperature. After incubation, TCEP was removed using a desalting column (PD-10 Sephadex G-25; Cytiva) equilibrated in SEC Buffer. The reduced ubiquitin was eluted from the desalting column directly into DBCO-maleimide (freshly resuspended in DMSO) such that there was a 5-fold molar excess of DBCO-Maleimide to cysteines, vortexed to mix, then incubated for 2 hours at room temperature. After incubation, excess (unbound) DBCO-maleimide was removed using a fresh desalting column. At this point, the ubiquitin with two DBCO “handles” (“DBCO-Ub”; DBCO-maleimide bound to cysteines) was combined with azide-ssDNA to form chimera (“ssDNA-Ub”). BCA assays were used to measure protein concentration in the presence or absence of ssDNA.

To covalently attach ubiquitin to ssDNA, DBCO-Ub was mixed with azide-ssDNA in a 1:1 molar ratio at 3 µM, then incubated at room temperature for 2 hours and concentrated using 3K MWCO PES protein concentrators (Pierce). Next, circular ssDNA-Ub product was purified by SEC (as for ubiquitin above) followed by incubation with functionalized agarose beads which sequester any remaining non-circular products by forming covalent bonds with exposed reactive groups. Specifically, DBCO agarose (Vector Labs) and maleimide agarose (Cube Biotech) slurries were mixed 1:1 in a 1.5 mL Eppendorf tube, washed 3x with SEC Buffer (spinning down gently and removing supernatant each round). Then, the post-SEC sample was added to the beads and tilt-stirred at 20°C for 2 hours before spinning down gently and collecting the supernatant.

#### Addition of complementary strand

A dsDNA-Ub sample was prepared by addition of 150 µM of ssDNA complementary strand to 150 µM of ssDNA-Ub, with formation of dsDNA confirmed by monitoring ^1^H NMR spectra in the DNA imino region (12-14 ppm ^1^H) to verify the formation of Watson-Crick hydrogen bonds.

#### ssDNA(s)-Ub chimera synthesis and purification

The linear ssDNA(s)-Ub control sample was made using short ssDNA (“ssDNA(s)”) modified to contain a single azide group on its 3’ end (12 nucleotides; /5Phos/TGGTGGAAGCTTG/3AzideN/; Integrated DNA Technologies). This short ssDNA was mixed with DBCO-Ub at an 8:1 molar ratio, incubated at room temperature for 2 hours, then purified by SEC.

#### Gel image processing

All gels were stained with either SYPRO Ruby (protein) or SYBR Gold (DNA). While those stains fluoresce (the brighter the band, the more protein, until high optical density darkens the bands again at high concentration), our protein ladder (NEB# P7706) does not (the darker the band, the more protein). To account for this and make gel images easier to interpret, we inverted the color scheme of all non-ladder lanes. In all figures, the ladder shown was run on the same gel as the lanes it is displayed next to.

### Nuclear Magnetic Resonance (NMR) spectroscopy

#### General acquisition parameters

Solution NMR data were acquired at 25-40°C using an 800 MHz Bruker Avance III spectrometer equipped with a ^1^H/^13^C/^15^N-optimized cryoprobe. Unless otherwise noted, all spectra were acquired with 150-350 µM protein in 50 mM potassium phosphate buffer (pH 5.8) with 7% D_2_O for signal lock. 5 mM MgCl_2_ was added to the dsDNA-Ub sample to stabilize the DNA duplex. Topspin 4.3.0 (Bruker) and NMRPipe (Delaglio et al., 1995) were used for data processing, with all NMRPipe processing carried out on the NMRbox server (Maciejewski et al., 2017). Analyses were performed using NMRFx Analyst version 11.4.24 (Norris et al., 2016; Johnson, 2018) and Poky software suite (Lee et al., 2021).

#### Backbone chemical shift assignments

Backbone chemical shift assignments were completed using Bruker triple-resonance pulse sequences, using samples with 150-350 µM U-[^13^C,^15^N] double-labelled protein in 50 mM potassium phosphate buffer (pH 5.8) with 7% D_2_O for signal lock.

For apo-Ub and ssDNA-Ub, HN(CA)CO, HNCO, CBCA(CO)NH and HNCACB spectra were collected at 25°C with 2048*80*128 complex points in the ^1^H, ^15^N, and ^13^C dimensions, respectively (Sattler, 1999). An additional HNCACB experiment was collected for ssDNA-Ub at 40°C with 2048*80*100 complex points in the ^1^H, ^15^N, and ^13^C dimensions, respectively, at 40°C to increase sensitivity and facilitate comparison with dsDNA-Ub spectra at higher temperatures. Backbone chemical shift assignments of dsDNA-Ub were based on HNCA, HN(CA)CO and HNCO collected at 40°C. For sites exhibiting multiple conformers (about 9 ssDNA-Ub residues with different ^1^H^N^ and ^15^N chemical shifts), we only assigned shifts of the most abundant state. All triple-resonance experiments were collected using non-uniform sampling (NUS) at a rate of 15% of complex points in the ^15^N and ^13^C dimensions and reconstructed using SMILE (Kazimierczuk and Orekhov, 2011). The assigned ^1^H, ^15^N, and ^13^C chemical shifts of apo-Ub, ssDNA-Ub and dsDNA-Ub have been deposited in the BMRB under accession numbers 53195, 53189, and 53191, respectively.

^1^H^N^-^1^H^N^ NOEs between amide protons in ssDNA-Ub were measured with an ^15^N-edited NOESY experiment with a mixing time of 125 ms recorded on a 300 µM U-[^13^C,^15^N] ssDNA-Ub sample at 40°C, acquired using 20% NUS in the indirectly-detected dimensions of a 2048*64*110 (^1^H^N^, ^15^N, ^1^H) complex point dataset. All NOEs discussed in the text were confirmed by verifying the presence of both of the two expected cross peaks, e.g. a ^1^H^N^ (residue A)-^1^H^N^ (residue B) NOE required observation of both the A(^1^H^N^)-B(^1^H^N^)-B(^15^N) and B(^1^H^N^)-A(^1^H^N^)-A(^15^N) peaks.

#### Calculation of chemical shift perturbations

Chemical shift perturbation calculations were performed using the following relationship:

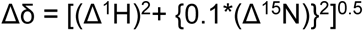

where the Δ^1^H and Δ^15^N denote the differences of the chemical shifts in ppm in the backbone amide proton and nitrogen dimension respectively, between samples as indicated. Comparisons to wild type ubiquitin (1–76) utilized prior assignments by (BioMagResBank accession number 5387) (Cornilescu et al., 1998).

#### Calculation of secondary structural propensities and order parameters

Using the backbone atom chemical shifts (^1^H^N^, ^15^N, ^13^Cα, ^13^Cβ, and ^13^CO for apo-Ub and ssDNA-Ub; ^1^H^N^, ^15^N, ^13^Cα, and ^13^CO for dsDNA-Ub), secondary structure propensities and order parameter values were calculated for all three constructs separately using MICS (Shen and Bax, 2012). All structure figures were generated with PyMOL (PyMOL Molecular Graphics System, version 3.5.1, Schrodinger, LLC, New York, NY).

Supplementary Information include Supplementary Figures: S1 – S10, Supplementary Tables: S1 – S2, and Supplementary References.

## Acknowledgements

We thank Denize C. Favaro for her outstanding support of the CUNY Advanced Science Research Center Biomolecular NMR Facility and assistance with aspects of this work, Dr. Giovanni Zocchi (UCLA) for insightful discussions, and SeArre Abebe (Harvard) for replicating the ubiquitin-DNA synthesis. This work was supported by NIH grants R01 GM106239 and R35 GM156269 (to KHG), DP2 GM141000 (to DRH), U54 CA132378 and U54 CA137788 (supporting SB); the Harvard College Research Program (to DG); Harvard Graduate School of Arts and Sciences (to MDS).

## Data Availability Statement

Backbone chemical shift assignments for apo-Ub, ssDNA-Ub, and dsDNA-Ub have been deposited in the Biological Magnetic Resonance Data Bank (bmrb.io) under accession numbers 53195, 53189, and 53191, respectively. Source data for gel images are available on Zenodo (DOI: 105281/zenodo.21072306).

## Author Contributions

DRH and KHG conceived the project. MDS devised the protein-DNA chimera synthesis workflow, and DG, MDS, and DRH developed the purification workflow. SB and KHG developed the NMR analysis strategy. MDS performed protein expression and purification, with early contributions from DG, and carried out the DBCO/azide conjugation chemistry, chimera purification, gel analyses, and NMR sample preparation. SB collected and assigned the NMR spectra. SB and KHG analyzed the chemical shift perturbations, MICS-derived secondary structure propensities and order parameters, and NOESY data. SB prepared and deposited the BMRB entries and organized the NMR source and supplementary data; MDS organized the remaining source and supplementary data. MDS and SB wrote the original draft, with MDS leading the protein-DNA chimera synthesis and purification components and SB leading the NMR spectroscopy and analysis components. All authors reviewed and edited the manuscript, with major editorial contributions from KHG and DRH. KHG supervised the CUNY/NMR component of the work, and DRH supervised the Harvard/protein-DNA chimera and mechanics component. KHG and DRH acquired funding.

## AI Disclosure Statement

The complete manuscript was first drafted without generative AI tools. ChatGPT was used during the proofreading stage to ensure consistency with journal formatting requirements, detect typographic and grammatical errors, and suggest language for the Significance and Author Contributions statements.

## Abbreviations

Apo-Ub: Ubiquitin bearing T9C, F45W, and K63C mutations
DBCO: dibenzo-cyclooctyne
DBCO-Ub: Apo-Ub with DBCO-maleimide attached at T9C and K63C (handles only)
ssDNA(s)-Ub: DBCO-Ub with short 12-nucleotide, single-stranded DNA individually attached to each DBCO group both at T9C and K63C
ssDNA-Ub: Apo-Ub with 50-nucleotide single-stranded DNA covalently linking T9C to K63C
dsDNA-Ub: ssDNA-Ub with its complementary DNA strand
NMR: Nuclear magnetic resonance
BMRB: Biological Magnetic Resonance Data Bank, https://bmrb.io
SEC: size exclusion chromatography

## Supplementary Information

**Figure S1:**
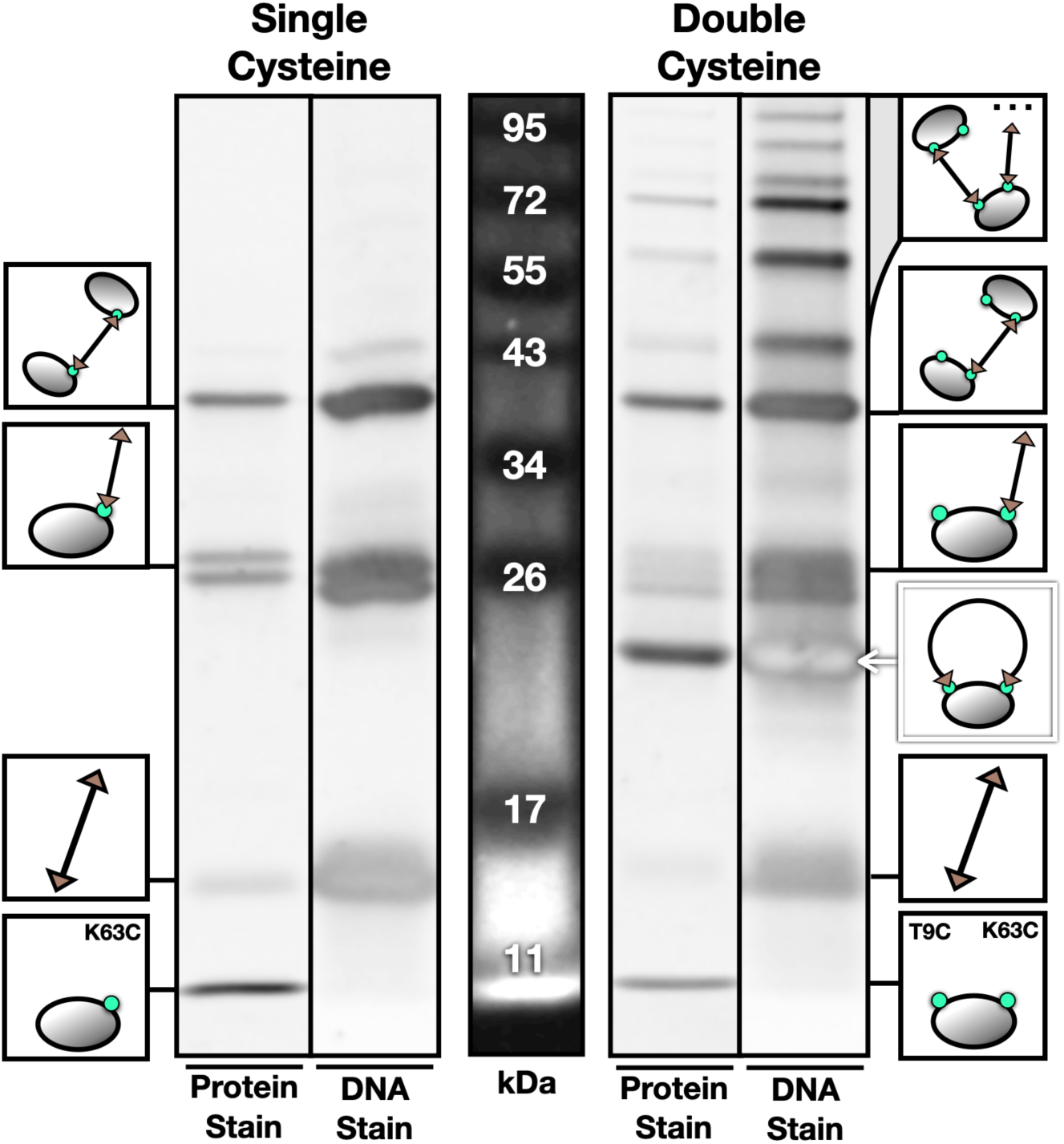
Protein and DNA Stains of Protein-ssDNA Products in SDS-PAGE. The same sample, stained for protein (SYPRO Ruby) and DNA (SYBR Gold) for single- and double-cysteine mutants, respectively. The DNA stain tends to be more sensitive and, at higher DNA concentrations, its intensity inverts (e.g. circular protein-ssDNA band). Trace amounts of DNA are stained by SYPRO Ruby (e.g. azide-ssDNA band).

**Figure S2:**
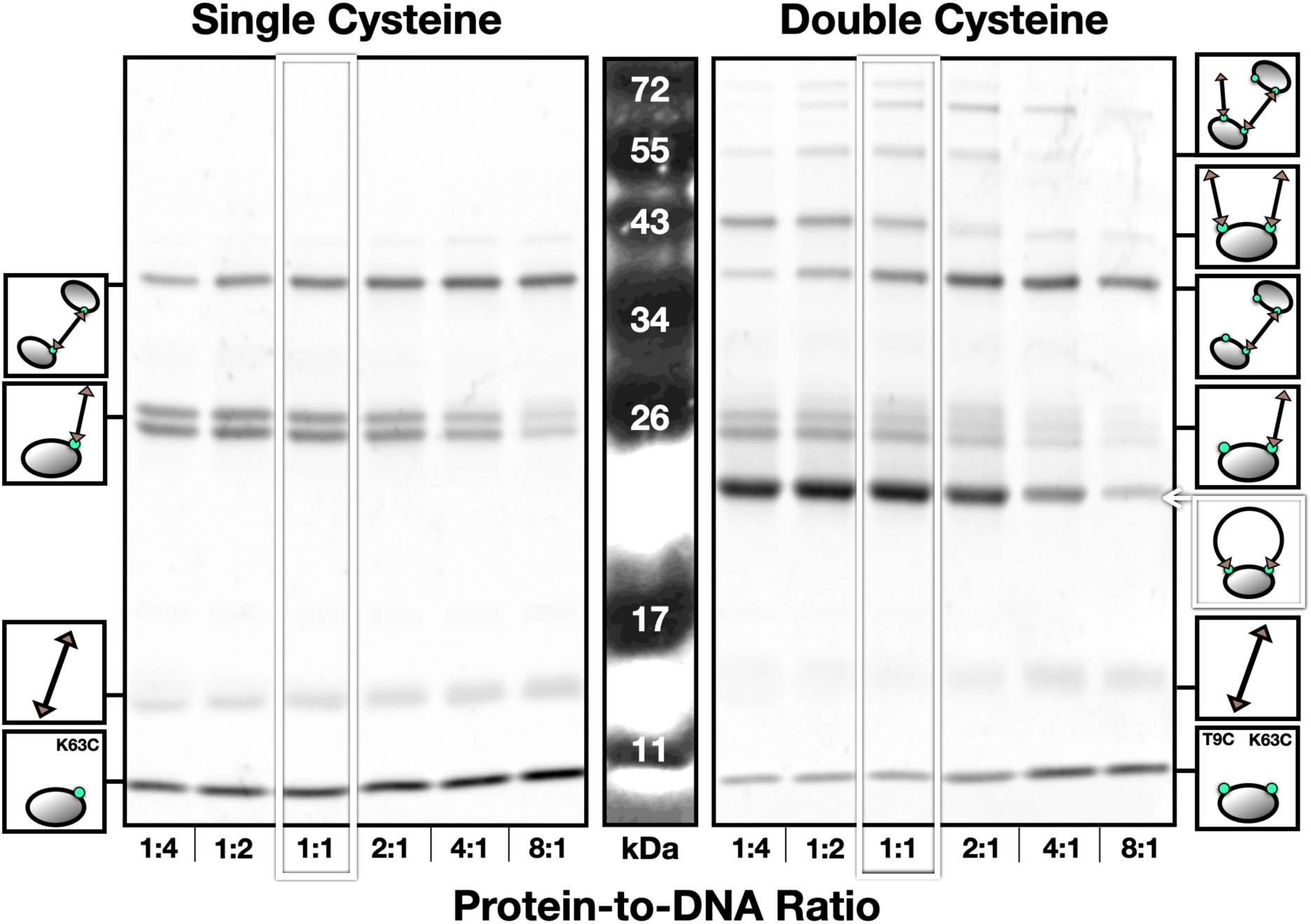
The Impact of Reactant Ratio on Synthesis Products. A higher protein:DNA ratio skews the mixture of products towards those with higher protein content. A ratio of ∼1:1 tends to be optimal for producing circular product. For these reactions, azide-ssDNA concentration was held constant (12 µM) while protein concentration was varied (3, 6, 12, 24, 48, 96 µM). Post-reaction, samples were normalized for protein concentration (all diluted to 3 µM protein), then immediately run on 12% SDS-PAGE at 100 V for 90 minutes. Gel stained for protein with SYPRO Ruby.

**Figure S3:**
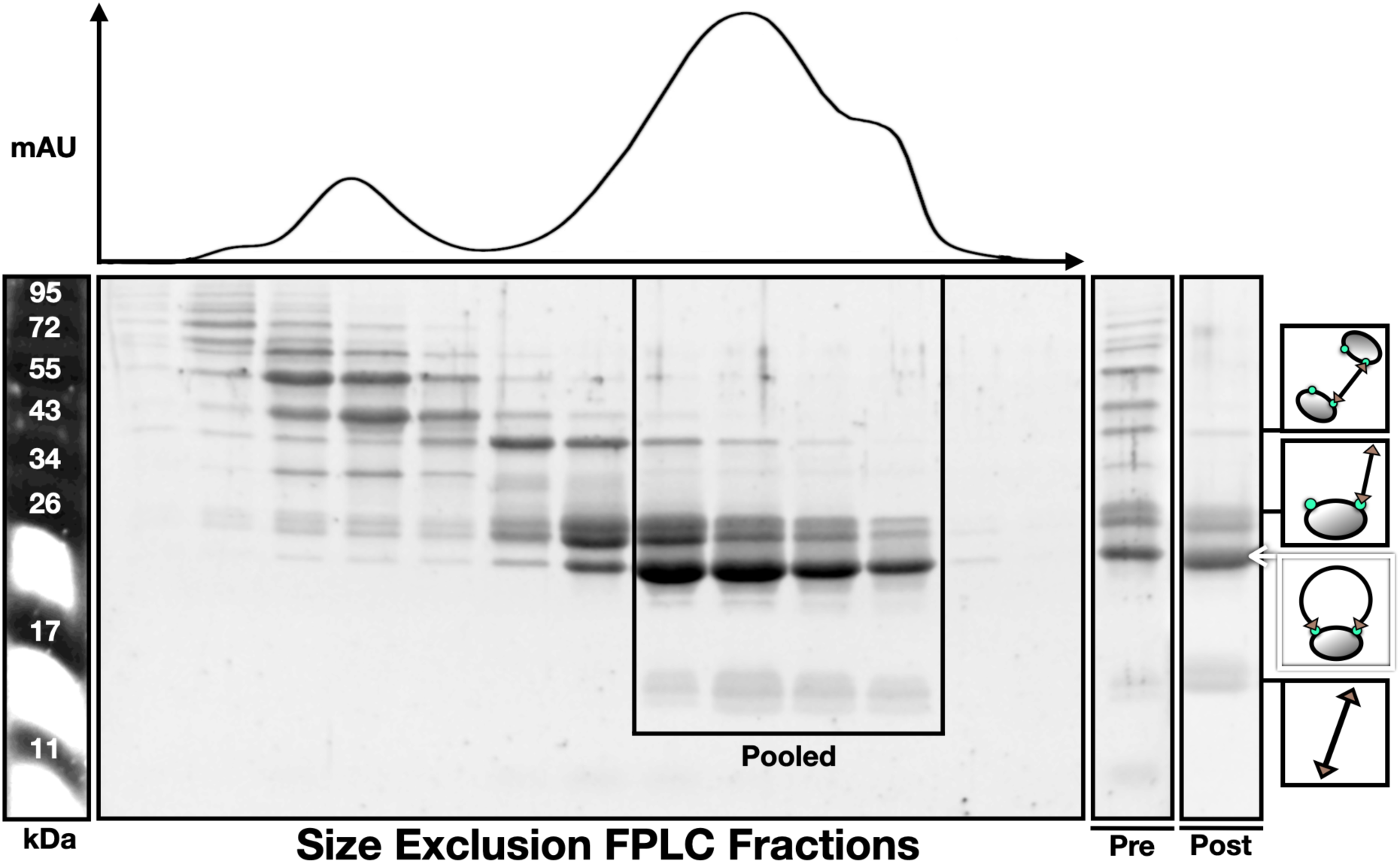
Size Exclusion Chromatography Purification of Circular Protein-ssDNA. Top: UV trace of size exclusion chromatography (SEC) using a Cytiva HiLoad Superdex 75 16/600 column. Bottom: native polyacrylamide gel electrophoresis of SEC fractions with Ubiquitin-ssDNA synthesis products. The indicated fractions are pooled and concentrated, increasing the concentration of circular product relative to other species (see “Pre” vs. “Post” on the right). Further purity can be achieved using reactive agarose, as shown in Figure 3.

**Figure S4:**
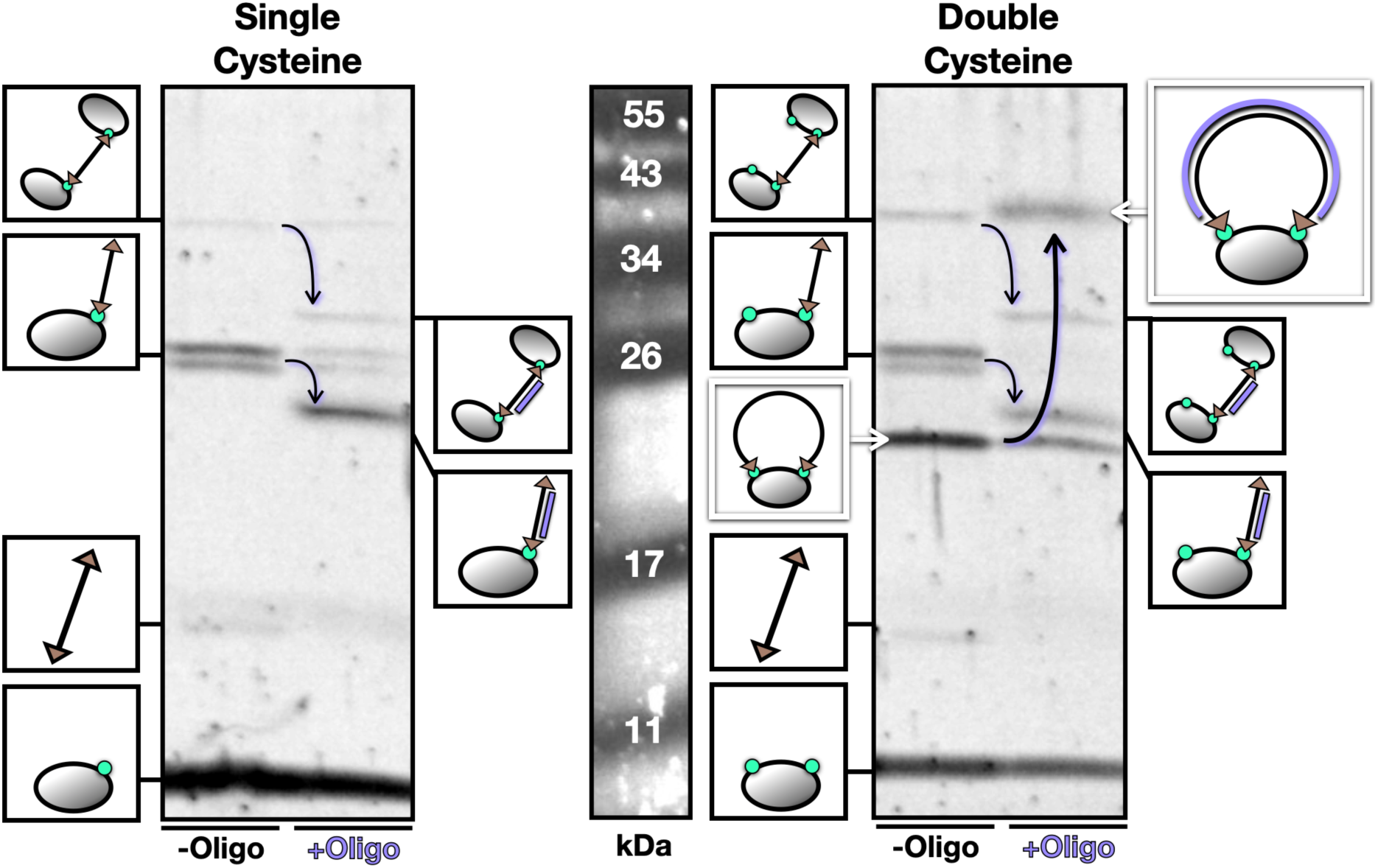
Identifying Protein-<u>ds</u>DNA Synthesis Products in SDS-PAGE. Addition of complementary ssDNA leads to formation of protein-dsDNA chimeras. The larger relative rigidity of double-stranded DNA causes products to migrate at unexpected speeds through the polyacrylamide gel, as previously observed (Gaillard & Strauss, 2015; Waszkiewicz et al., 2023). Here, addition of non-saturating quantities of complementary strand (∼1:2 molar ratio) results in incomplete band shifts. For complete band shift (∼1.2:1 molar ratio), see Figure 4.

**Figure S5:**
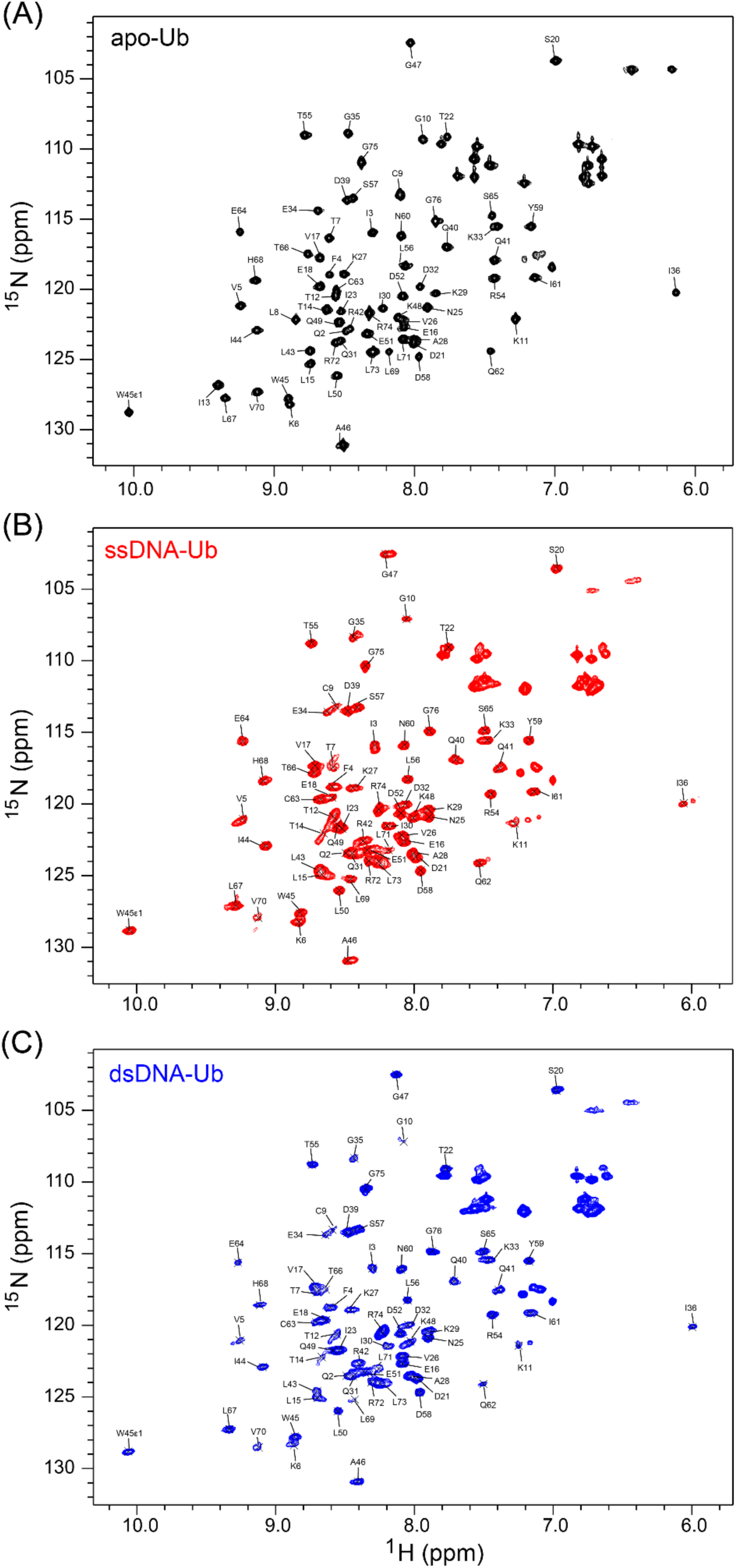
^15^N-^1^H HSQC Spectra Of Ubiquitin And Ubiquitin-DNA Conjugates, With Residue Assignments At 40°C. (A) apo–Ub, (B) ssDNA-Ub and (C) dsDNA-Ub.

**Figure S6:**
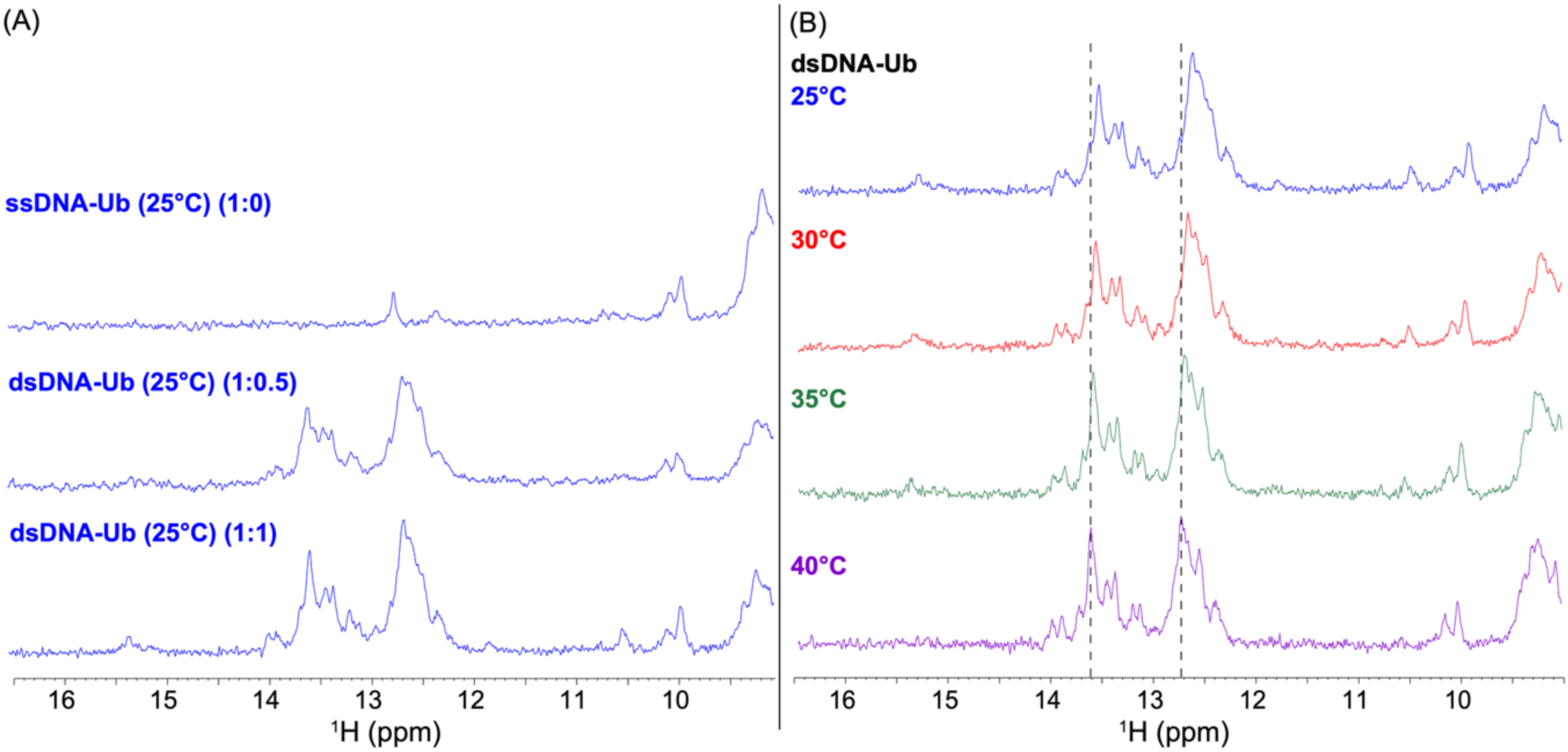
DNA Imino ^1^H NMR Signals Confirm The Formation Of Watson-Crick Base Pairing With Double Stranded DNA. (A) Downfield region of ^1^H NMR spectrum, including DNA imino ^1^H signals expected downfield of 12 ppm. Spectra obtained from 150 µM ssDNA-Ub (top) and dsDNA-Ub (middle with 0.5 equivalents of complementary strand, bottom with 1.0 equivalent), all at 25°C. (B) Same region of ^1^H NMR spectrum, with multiple spectra collected on 150 µM dsDNA-Ub between 25°C and 40°C as indicated. Imino ^1^H signals sharpen and shift downfield with increasing temperature; dashed lines provide a vertical guide to the eye.

**Figure S7:**
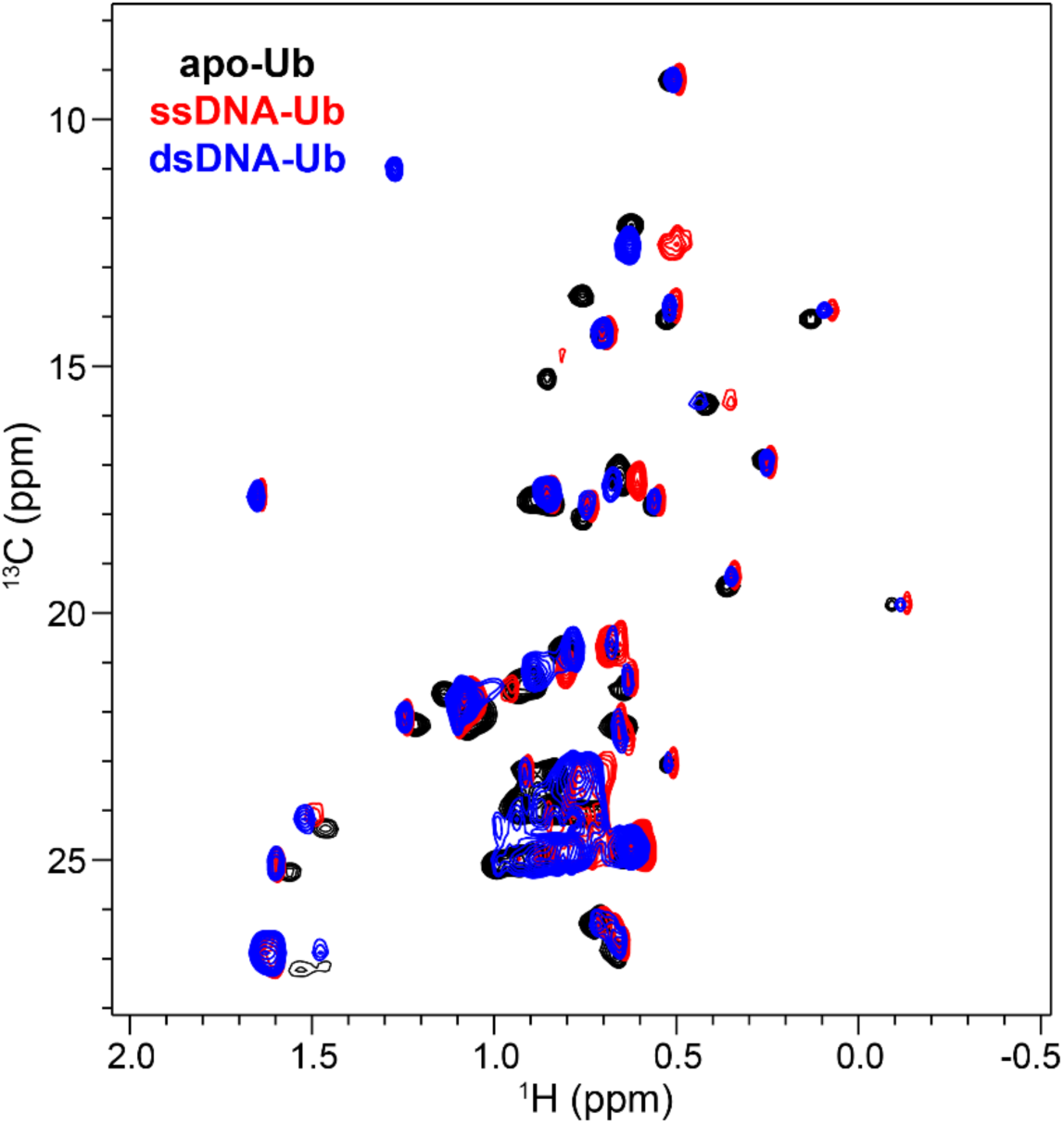
Overlaid Methyl Regions Of Constant-Time ^13^C-^1^H HSQCs of apo-Ub (black), ssDNA-Ub (red), and dsDNA-Ub (blue) at 40°C.

**Figure S8:**
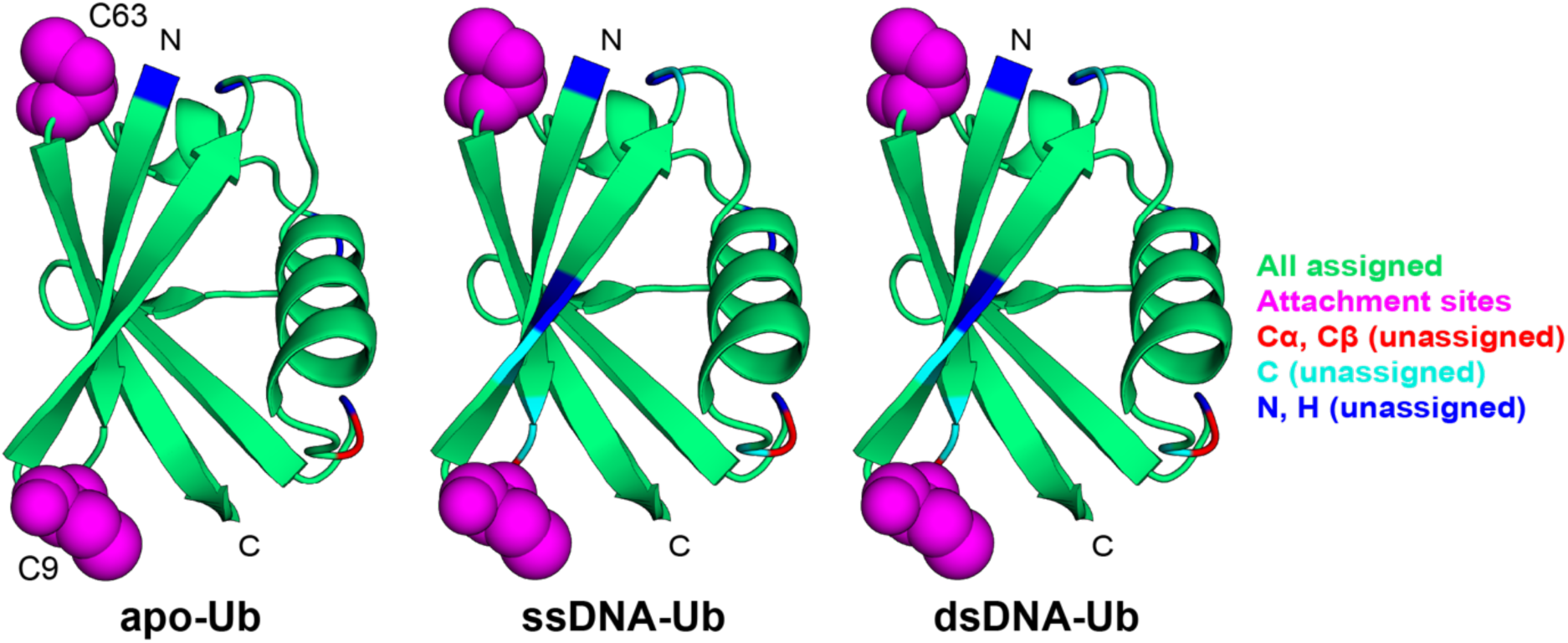
Ribbon Diagram Of Ubiquitin (PDB: 1UBQ) Showing High Degree Of Assignment Completeness. Color codes are indicated on the right side. The N- and C-termini are indicated, as are the locations of the T9C and K63C point mutations made to establish the covalent attachment points.

**Figure S9:**
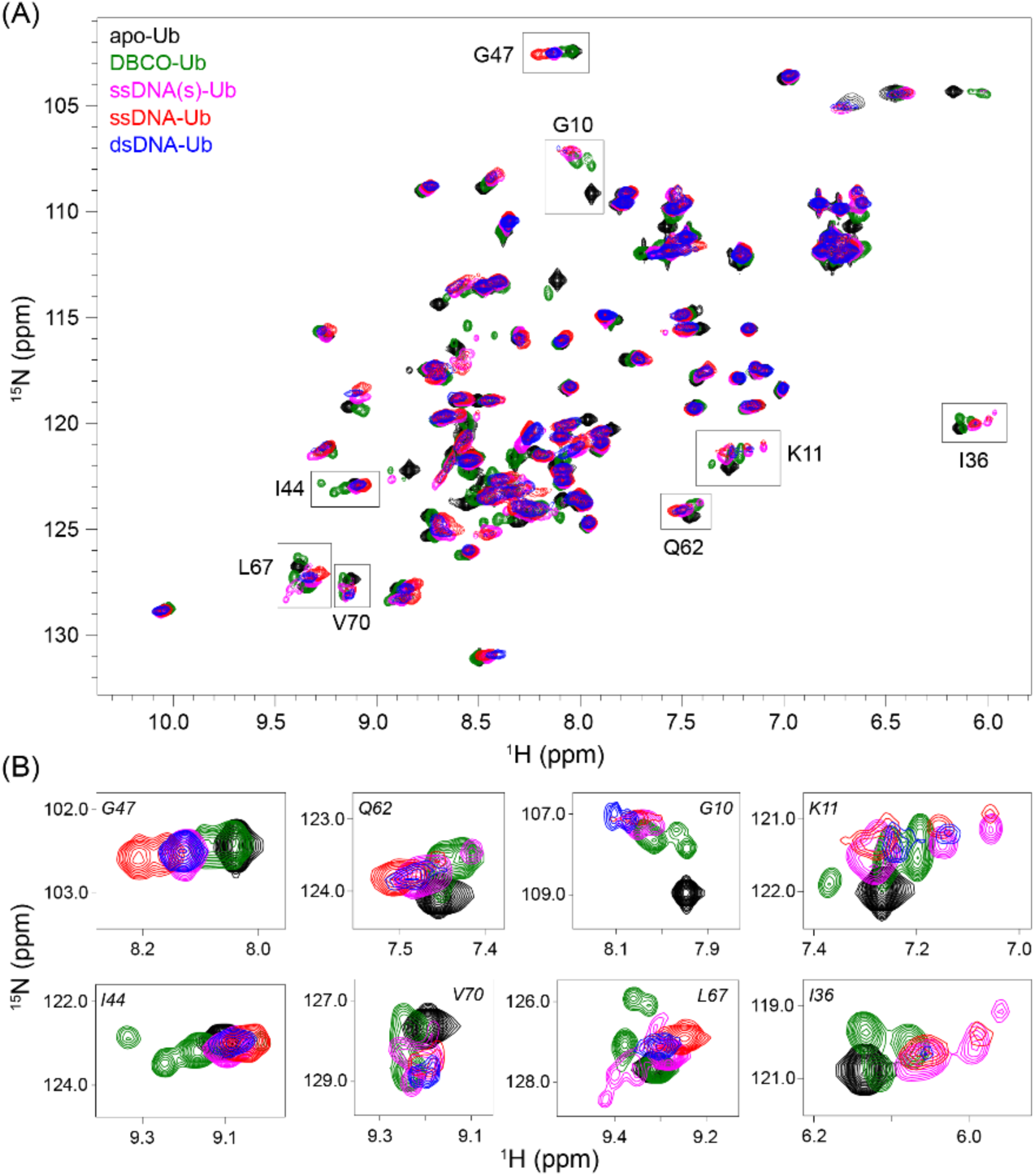
DBCO Tethering Induces Conformational Heterogeneity In Ub, Subsequently Reduced By Addition Of Short 12-nt ssDNA. (A) Overlay of ^15^N/^1^H HSQC spectra of apo-Ub (black), DBCO-Ub (green), ssDNA(s)-Ub (magenta), ssDNA-Ub (red), and dsDNA-Ub (blue) at 40°C at pH 5.8. Residues showing conformational heterogeneity are indicated with boxes. (B) Overlay of ^15^N/^1^H HSQC spectra for selected resonances, using same spectra as (A).

**Figure S10:**
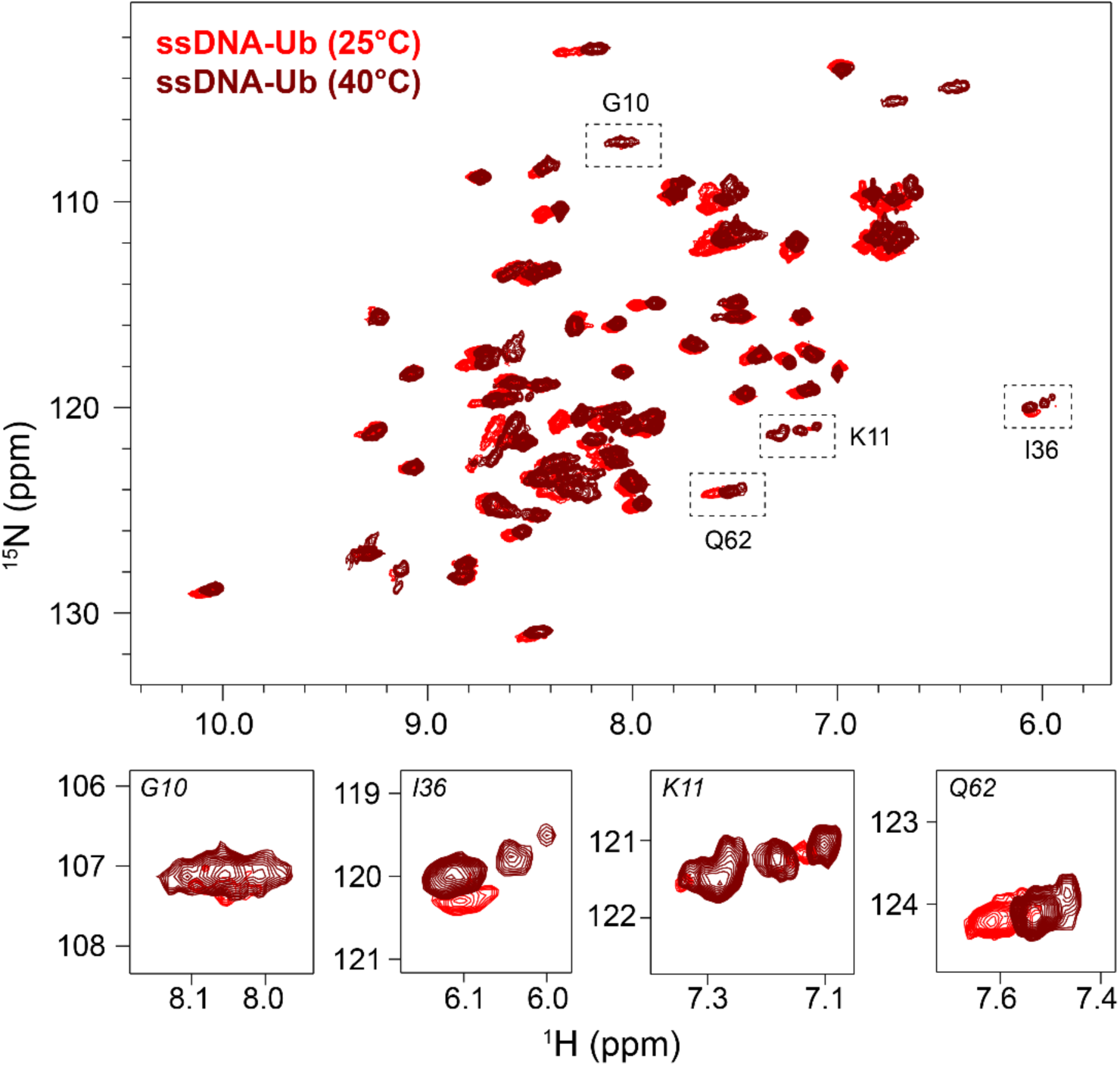
Overlaid ^15^N-^1^H HSQC Spectra Of ssDNA-Ub At 25°C Or 40°C. Residues exhibiting temperature-dependent changes showed altered relative populations of heterogeneous resonances. Regions with temperature-induced conformational shifts are highlighted with dotted boxes (top) and displayed as enlarged views (bottom).

**Table S1:** Assignment completeness (%) of three different constructs:

| Name of atom | Completeness (%) |  |  |
| --- | --- | --- | --- |
|  | Apo-Ub | ssDNA-Ub | dsDNA-Ub |
| Cα | 75/76 = 98.68 | 73/76 = 96.05 | 73/76 = 96.05 |
| Cβ | 69/70 = 98.57 | 65/70 = 92.85 | N.A. |
| C | 75/76 = 98.68 | 69/76 = 90.78 | 69/76 = 90.78 |
| N | 70/73 = 95.89 | 68/73 = 93.15 | 68/73 = 93.15 |
| H | 70/73 = 95.89 | 68/73 = 93.15 | 68/73 = 93.15 |
| Total | 359/368 = 97.6 | 343/368 = 93.21 | 278/298 = 93.29 |

(N.A.: Not applicable, as Cβ assignments were not collected on dsDNA-Ub)

**Table S2:** List of ^1^H^N^-^1^H^N^ NOEs between β5-β1 and β5-β3 strands for wild-type Ub (1UBQ) and slipped Ub (5OXI) conformers. The predicted NOEs are shown in plain font, predicted and observed NOEs are shown in *italics*, and newly observed NOEs are shown in **bold**.

| $\beta 3$ | $\beta 5$ | $\beta 1$ | Distance (Å)<br>1UBQ | Distance (Å)<br>5OXI |
| --- | --- | --- | --- | --- |
|  | 64 | 2 | 3.9 | 9.3 |
|  | 65 | 4 | 4.4 | 9.7 |
|  | 66 | 4 | 4.7 | 8.3 |
|  | 67 | 5 | 4.7 | 8.7 |
|  | 67 | 6 | 3.6 | 9.9 |
|  | 68 | 4 | 8.3 | 5 |
|  | 68 | 6 | 4.9 | 7.8 |
|  | 69 | 5 | 7.3 | 4.6 |
|  | 69 | 6 | 3.8 | 3.7 |
|  | 70 | 6 | 7.9 | 4.7 |
| 44 | 68 |  | 2.8 | 10.2 |
| 44 | 69 |  | 4.9 | 6.9 |
| 42 | 70 |  | 3.1 | 8.6 |
| 43 | 70 |  | 5 | 6.9 |
| 45 | 70 |  | 8.6 | 5.1 |
| 44 | 70 |  | 4.5 | 2.7 |
|  | 71 | 6 | 10.6 | 3.9 |
|  | 71 | 7 | 9.2 | 5.1 |
| 41 | 72 |  | 4.8 | 7.3 |
| 40 | 72 |  | 5.5 | 9.3 |
| 42 | 72 |  | 4.4 | 2.9 |
| 43 | 72 |  | 8.2 | 5 |
| 42 | 73 |  | 8.3 | 5.1 |
| **40** | **73** |  | **8.2** | **9.5** |
| **42** | **69** |  | **6.9** | **12.1** |
|  | **65** | **3** | **6.1** | **11.3** |

## References

Bustamante C, Yan S. The development of single molecule force spectroscopy: from polymer biophysics to molecular machines. Q Rev Biophys. 2022; 55:e9. 10.1017/S0033583522000087

Chen H, Meisburger SP, Pabit SA, Sutton JL, Webb WW, Pollack L. Ionic strength-dependent persistence lengths of single-stranded RNA and DNA. Proc Natl Acad Sci U S A. 2012; 109:799–804. 10.1073/pnas.1119057109

Cornilescu G, Marquardt JL, Ottiger M, Bax A. Validation of Protein Structure from Anisotropic Carbonyl Chemical Shifts in a Dilute Liquid Crystalline Phase. Journal of the American Chemical Society. 1998; 120:6836–6837. 10.1021/ja9812610

Delaglio F, Grzesiek S, Vuister G, Zhu G, Pfeifer J, Bax A. NMRPipe: a multidimensional spectral processing system based on UNIX pipes. Journal of biomolecular NMR. 1995; 6:277–293. 10.1007/BF00197809

Driscoll P, Thompson g, Harris R. The Ubiquitin NMR Resource - Archive 1 [Backbone Experiments]. [Data set] In The Ubiquitin NMR Resource. 2025; 969:114–126. 10.5281/zenodo.14791182

Gaillard C, Strauss F. Construction of DNA Hemicatenanes from Two Small Circular DNA Molecules. PLOS ONE. 2015; 10:e0119368. 10.1371/journal.pone.0119368

Gladkova C, Schubert AF, Wagstaff JL, Pruneda JN, Freund SMV, Komander D. An invisible ubiquitin conformation is required for efficient phosphorylation by PINK1. The EMBO Journal. 2017; 36:3555–3572. 10.15252/embj.201797876

Gokulu IS, Banta S. Enzyme Engineering by Force: DNA Springs for the Modulation of Biocatalytic Trajectories. ACS Synth Biol. 2024; 13:2600–2610. 10.1021/acssynbio.4c00431

Guilbaud S, Salome L, Destainville N, Manghi M, Tardin C. Dependence of DNA Persistence Length on Ionic Strength and Ion Type. Physical Review Letters. 2019; 122:028102. 10.1103/PhysRevLett.122.028102

Johnson BA. From Raw Data to Protein Backbone Chemical Shifts Using NMRFx Processi. Methods in Molecular Biology. 2018. 10.1007/978-1-4939-7386-6_13

Joseph C, Tseng CY, Zocchi G, Tlusty T. Asymmetric effect of mechanical stress on the forward and reverse reaction catalyzed by an enzyme. PLoS One. 2014; 9:e101442. 10.1371/journal.pone.0101442

Kay LE. New Views of Functionally Dynamic Proteins by Solution NMR Spectroscopy. J Mol Biol. 2016; 428:323–331. 10.1016/j.jmb.2015.11.028

Kazimierczuk K, Orekhov VY. Accelerated NMR Spectroscopy by Using Compressed Sensing. Angewandte Chemie International Edition. 2011; 50:5556–5559. 10.1002/anie.201100370

Khorasanizadeh S, Peters ID, Butt TR, Roder H. Folding and stability of a tryptophan-containing mutant of ubiquitin. Biochemistry. 1993; 32:7054–7063. 10.1021/bi00078a034

Kim Y, Prestegard JH. A dynamic model for the structure of acyl carrier protein in solution. Biochemistry. 1989; 28. 10.1021/bi00448a017

Komander D, Rape M. The Ubiquitin Code. Annual Review of Biochemistry. 2012; 81:203–229. 10.1146/annurev-biochem-060310-170328

Lee W, Rahimi M, Lee Y, Chiu A. POKY: a software suite for multidimensional NMR and 3D structure calculation of biomolecules. Bioinformatics (Oxford, England). 2021:1–2. 10.1093/bioinformatics/btab180

Maciejewski MW, Schuyler AD, Gryk MR, Moraru II, Romero PR, Ulrich EL, Eghbalnia HR, Livny M, Delaglio F, Hoch JC. NMRbox: A Resource for Biomolecular NMR Computation. Biophysical Journal. 2017; 112. 10.1016/j.bpj.2017.03.011

Mukhortava A, Schlierf M. Correction to Efficient Formation of Site-Specific Protein–DNA Hybrids Using Copper-Free Click Chemistry. Bioconjugate Chemistry. 2016; 27:1559–1563. 10.1021/acs.bioconjchem.6b00361

Norris M, Fetler B, Marchant J, Johnson BA. NMRFx Processor: a cross-platform NMR data processing program. Journal of Biomolecular NMR. 2016; 65:205–216. 10.1007/s10858-016-0049-6

Pickart CM, Raasi S. Controlled Synthesis of Polyubiquitin Chains. Methods in Enzymology. 2005; 399:21–36. 10.1016/S0076-6879(05)99002-2

Sattler M. Heteronuclear multidimensional NMR experiments for the structure determination of proteins in solution employing pulsed field gradients. Progress in Nuclear Magnetic Resonance Spectroscopy. 1999; 34:93–158. 10.1016/S0079-6565(98)00025-9

Schubert AF, Gladkova C, Pardon E, Wagstaff JL, Freund SMV, Steyaert J, Maslen SL, Komander D. Structure of PINK1 in complex with its substrate ubiquitin. Nature. 2017; 552:51–56. 10.1038/nature24645

Shen Y, Bax A. Identification of helix capping and β-turn motifs from NMR chemical shifts. Journal of Biomolecular NMR. 2012; 52:211–232. 10.1007/s10858-012-9602-0

Tseng CY, Wang Y, Zocchi G. Enzyme-DNA chimeras: Construction, allostery, applications. Methods Enzymol. 2021; 647:257–281. 10.1016/bs.mie.2020.09.010

Tseng CY, Zocchi G. Mechanical control of Renilla luciferase. J Am Chem Soc. 2013; 135:11879–11886. 10.1021/ja4043565

Vijay-kumar S, Bugg CE, Cook WJ. Structure of ubiquitin refined at 1.8 Å resolution. Journal of Molecular Biology. 1987; 194:531–544. 10.1016/0022-2836(87)90679-6

Wang A, Zocchi G. Elastic energy driven polymerization. Biophys J. 2009; 96:2344–2352. 10.1016/j.bpj.2008.11.065

Wang Y, Wang A, Qu H, Zocchi G. Protein-DNA chimeras: synthesis of two-arm chimeras and non-mechanical effects of the DNA spring. J Phys Condens Matter. 2009; 21:335103. 10.1088/0953-8984/21/33/335103

Waszkiewicz R, Ranasinghe M, Fogg JM, Catanese DJ, Ekiel-Jeżewska ML, Lisicki M, Demeler B, Zechiedrich L, Szymczak P. DNA supercoiling-induced shapes alter minicircle hydrodynamic properties. Nucleic Acids Research. 2023; 51:4027–4042. 10.1093/nar/gkad183

Wauer T, Simicek M, Schubert A, Komander D. Mechanism of phospho-ubiquitin-induced PARKIN activation. Nature. 2015; 524:370–374. 10.1038/nature14879

Zocchi G. Molecular Machines: A Materials Science Approach. Princeton, NJ: Princeton University Press. 2018. 10.23943/9781400890064

## Supplementary References

Gaillard C, Strauss F (2015). Construction of DNA Hemicatenanes from Two Small Circular DNA Molecules. PLOS ONE 10, e0119368. 10.1371/journal.pone.0119368.

Waszkiewicz R, Ranasinghe M, Fogg JM, Catanese DJ, Ekiel-Jeżewska ML, Lisicki M, Demeler B, Zechiedrich L, Szymczak P (2023). DNA supercoiling-induced shapes alter minicircle hydrodynamic properties. Nucleic Acids Research 51, 4027–4042. 10.1093/nar/gkad183.

